# Tokenizing Foldable Protein Structures with Machine-Learned Artificial Amino-Acid Vocabulary

**DOI:** 10.1101/2023.11.27.568722

**Authors:** Xiaohan Lin, Zhenyu Chen, Yanheng Li, Zicheng Ma, Chuanliu Fan, Ziqiang Cao, Shihao Feng, Yi Qin Gao, Jun Zhang

## Abstract

Understanding functions of proteins and designing proteins committed to specific functions *in silico* are highly valuable for science, industry and therapeutics. However, there is a long-standing divergence in how to present function-related protein structures to the machine learning models: Although the 1-dimensional (1D) representation of proteins via Anfinsen’s tokens (i.e., amino acids) is sufficient in principle and more machine-friendly, it is less successful in structure-oriented protein design compared to symmetry-constrained 3D representation (i.e., atom coordinates). Aiming to bridge the gap between 1D and 3D protein representations and harvest the advantages of the two, we develop *probabilistic tokenization theory* for metastable protein structures. We present an unsupervised learning strategy, which conjugates inverse folding with structure prediction, to encode protein structures into artificial amino-acid tokens (ProTokens) and decode them back to atom coordinates. We show that tokenizing protein structures *variationally* can lead to compact and informative representations. Compared to amino acids — the Anfinsen’s tokens — ProTokens are easier to detokenize and more descriptive of finer conformational ensembles. Therefore, protein structures can be efficiently compressed, stored, aligned and compared in the form of ProTokens. By unifying the discrete and continuous representations of protein structures, ProTokens also enable all-atom protein structure design via various generative models without the concern of symmetry or modality mismatch, and allows scalable foundation models to perceive, process and explore the microscopic structures of biomolecules effectively.

## 1. Introduction

Understanding functions of proteins and designing proteins committed to specific functions are highly valuable in science, industry and therapeutics. Considering that a protein’s function is largely determined by its 3D structure, recently, *in silico* (particularly, AI-aided) parse and design of protein structures are gaining increasing attention. However, there is a long-standing debate in how to present function-related protein structures to the machine learning models, causing divergence in the ongoing research momentum. Based on Anfinsen’s hypothesis, one popular strategy is to represent a protein structure by its amino acid sequence (i.e., 1D representation), which is simple and AI-friendly, allowing generation of all-atom protein structures with off-the-shelf AI models. Unfortunately, the translation from the generated amino acids into real 3D atom coordinates remains a non-trivial task. In addition, alternative protein conformations corresponding to different functions are indistinguishable in terms of 1D representation. These limitations motivate an orthogonal approach for protein structure research, where the structures are represented by the spatial coordinates of protein atoms (3D representation). However, protein 3D structures are defined by spatial coordinates of atoms which exhibit specific symmetry and are subjected to physics constraints (such as transrotational equivariance and polymer restraints), hence, significantly impeding the transaction of modern generative artificial intelligence (AI), e.g., generative adversarial(Goodfellow et al., 2014), diffusion(Ho et al., 2020; Song et al., 2021; Song and Ermon, 2019) and autoregressive(Chen et al., 2017; Papamakarios et al., 2017; Salimans et al., 2017) models etc., for protein design. Up to now, 3D representation still encounters significant difficulty of being compatible with scalable AI models due to the symmetry issue.(Watson et al., 2023) Even if a symmetry-constraint AI is deliberately designed, joint generation of all-atom structures (i.e., backbones and sidechains) as in 1D representation remains unresolved.(Ingraham et al., 2023; Jing et al., 2023)

On the other hand, with the rise of large language models (LLMs) as a potential artificial general intelligence (AGI) candidate, many efforts are being paid to “unify the modality” of various signals (including images, videos and sounds etc.) to texts or tokens that LLMs are familiar with(Banerjee and Arora, 2023; Dosovitskiy et al., 2021; Ryoo et al., 2021). It is highly desired that a representation of proteins amenable to LLMs can be developed in order to give AGI an access to the microscopic biomolecular universe. But it is unwise to treat 3D coordinates of proteins directly as input or output of an LLM due to the symmetry issue. Indeed, the Cartesian atom coordinates of proteins are known to be redundant for representing protein structures given the trans-rotational equivariance and the polymer nature of protein chains. Consequently, the modality difference of protein structure representations (1D v.s. 3D) causes significant divergence in the research paradigms of proteins, particularly in the realm of protein design. In this research, we provide a unified perspective for protein 1D and 3D structures based on protein physics, with an important conclusion that although protein 3D structures are continuous, the function-related conformational ensembles (or metastable states) are countable at a proper observation timescale. Particularly, the conventionally defined 1D structure of proteins (i.e., amino acid sequence) is equivalent to a probabilistic tokenization of protein 3D structure ensemble given a large observation timescale when all the (un-)folded structures collapse to a single metastable state. Although being compact, amino acid tokens bear some shortcomings in terms of informativeness, including the difficulty of being tokenized to 3D structures and that alternative conformations corresponding to different functions become degenerate.

In this research, starting from the *Anfinsen’s tokens* (i.e., the set of amino acids), we attempt to create an artificial amino-acid vocabulary for protein 3D structures, trying to strike better balance between the compactness and informativeness of these probabilistic tokens. The main features of this study are multi-fold: (1) We propose and justify the probabilistic tokenization theory of protein structures, by factoring the continuous distribution of spatial coordinates into discrete parts representing function-relevant metastable states and continuous parts accounting for the conformational fluctuations within the metastable state. (2) We develop a data-driven and physics-informed approach to extracting amino-acid-like ProTokens in an unsupervised and variational way. The ProTokens are compact and informative representations for all-atom protein structure ensembles. (3) We unify the 1D and 3D modality of protein structures by merging amino acids as subset of ProToken vocabulary, and equip ProTokens with *Janus* representations, making them ready for both discrete language-based and continuous diffusion-based foundation models. (4) We summarize caveats and pitfalls of applying a transformed representation like ProTokens in tasks related to protein structures and develop mathematical guidelines to ameliorate these issues.

## 2. Results

### A. Functional protein structures can be reasonably tokenized via deep learning

#### 1. Probabilistic tokenization theory justifies physics-informed discretization of protein 3D structures and extends the vocabulary of amino acids

Protein 3D structures are usually presented as the spatial coordinates of the atoms which belong to the SE(3)-symmetry group. Such symmetry suggests it may not be optimal to treat 3D coordinates of proteins directly as input or output of any symmetry-free models including the state-of-the-art foundation models(Liu et al., 2024; OpenAI et al., 2024; Peebles and Xie, 2023). Furthermore, protein is a special kind of biopolymer, and its structure is subjected to physics restraints such as peptide bonding interactions which are usually not relevant to conformational changes or its functions. Consequently, the spatial coordinates of protein structure are a redundant representation and not amenable to modern AI. However, We note that, although the structure of a protein can be continuously changed in the Cartesian coordinate space, the *set of its metastable states* are countable, hence, can be well-defined discretely according to the landscape theory(Eaton et al., 2000; Ghosh and Ranjan, 2020). Specifically, the definition of metastable states depends on the observation timescale *τ*_obs_: A metastable state can be only defined if *τ*_rlx_ < *τ*_obs_ <*τ*_life_. According to the landscape theory(Konovalov et al., 2021; Wales, 2004), the smaller *τ*_obs_ is, the larger amount of metastable states can be defined(“Markov state models of biomolecular conformational dynamics,” 2014; Zhang et al., 2019) (Fig. 1a). In a limit case, when *τ*_obs_ is comparable to the (un-)folding timescale (*τ*_fold_) of a protein, most proteins are known to exhibit a two-state kinetics between the folded and unfolded states, hence, the only folded metastable state can be well defined by the amino-acid sequence of the protein (Fig. 1a).

**Fig. 1.**
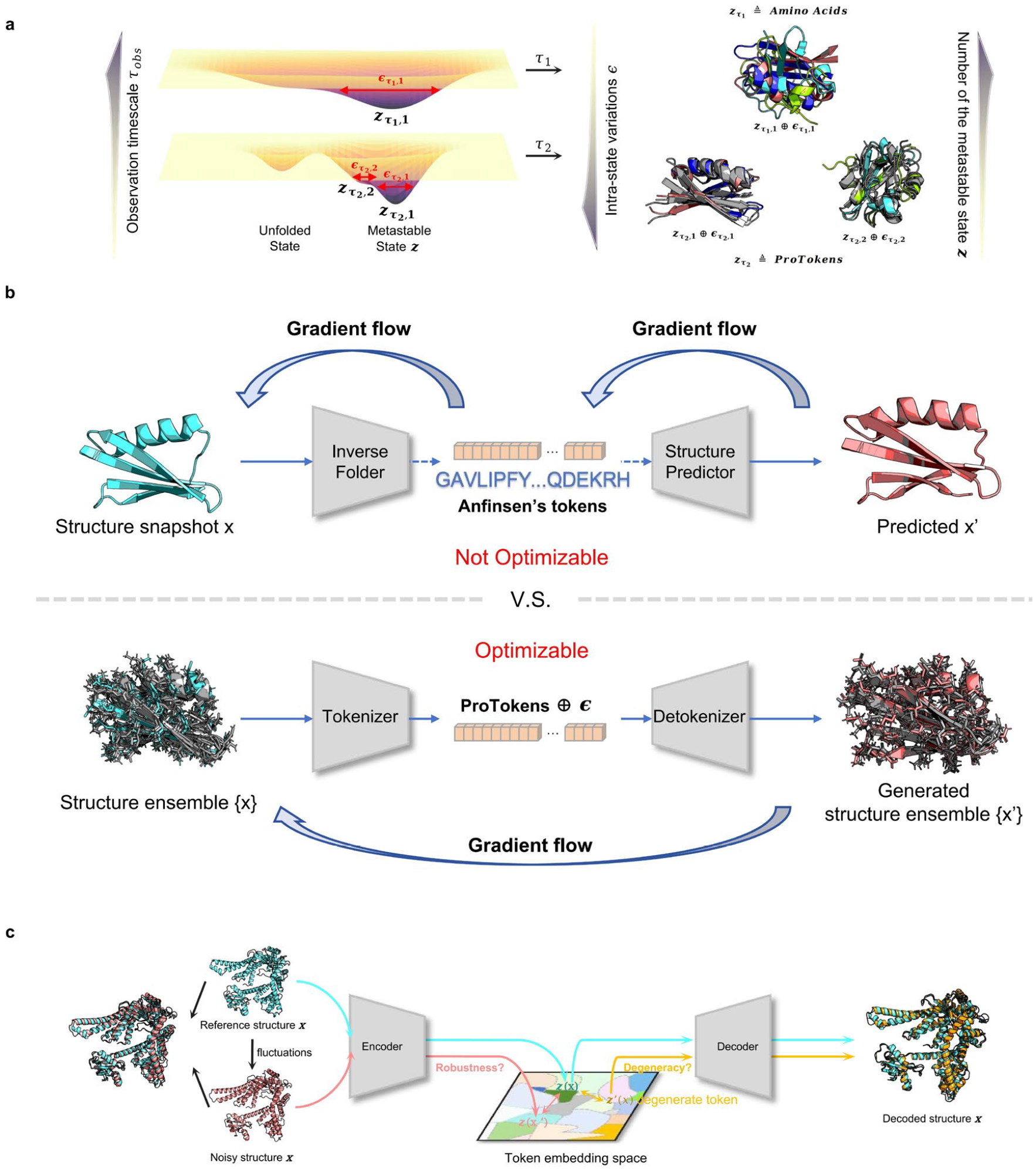
Probabilistic tokenization of protein 3D structures. **a**, Metastable states defined at different timescales can lead to different tokens of protein structures, including amino acids. **b**, ProTokens can be learned by connecting inverse folding and structure prediction models end-to-end, and replacing amino acids by optimizable vocabulary. **c**, Machine-learned tokens may suffer from non-robustness (red) and degeneracy (yellow).

**Fig. 2.**
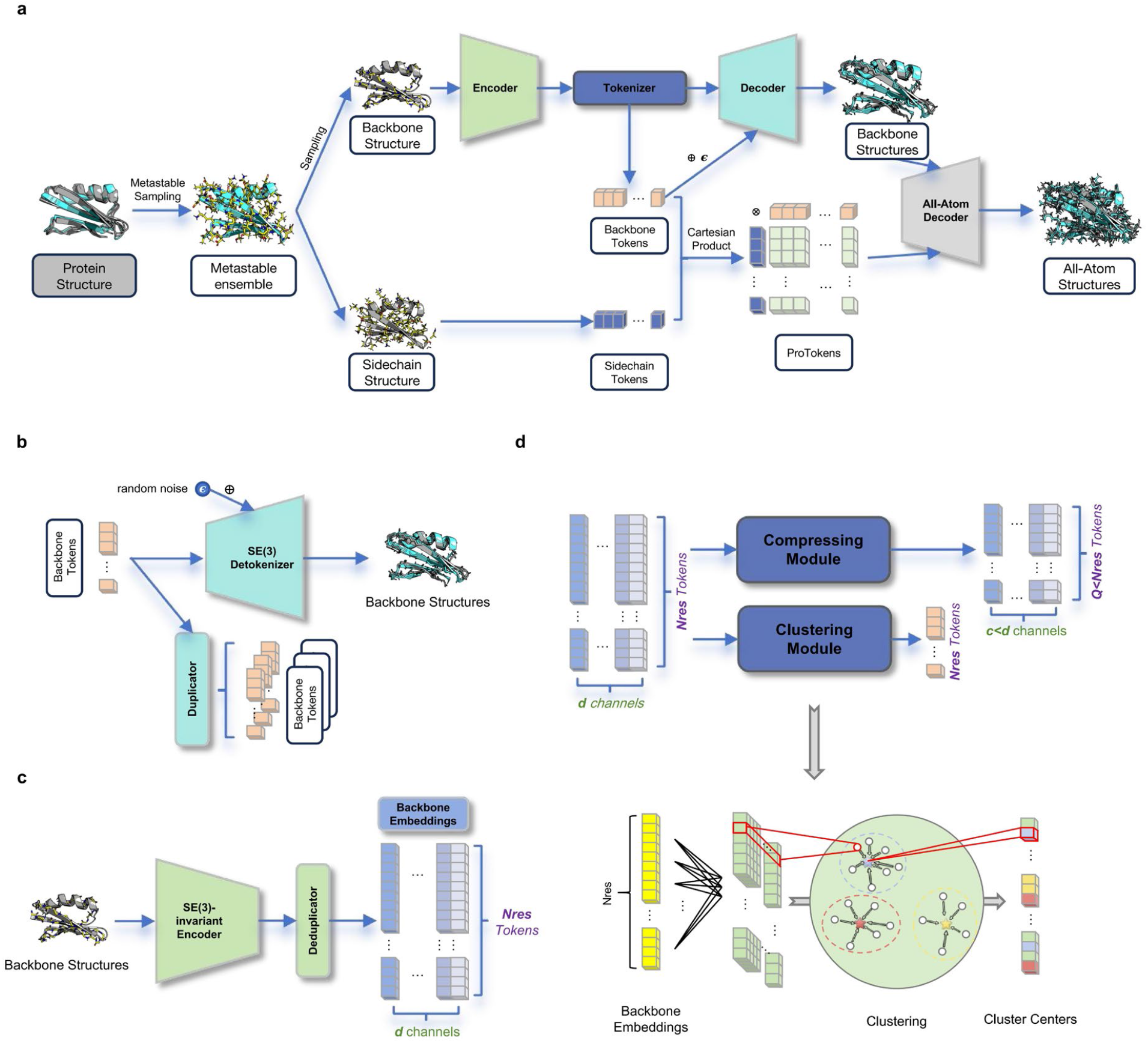
Architecture of ProToken Distiller. **a**, Overview of the ProToken Distiller. The all-atom structure(s) is tokenized through the backbone track and sidechain track separately, leading to backbone tokens and side chain tokens, respectively. ProToken is the Cartesian product of the two. ProToken Distiller consists of a generative Decoder **(b)**, an Encoder **(c)**, and a Tokenzier **(d)**.

Thanks to the metastability, a continuous distribution of function-related protein structures can now be reasonably factored into discrete and continuous parts,

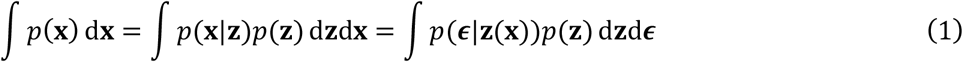

where the discrete part **z** stands for the metastable state, *p* (**z**) is a discrete distribution specifying the number of metastable states which **x**∼*p* (**x**) consists of, and the continuous part *p***(ϵ**|**z(x))** covers the intrinsic structure fluctuations within this metastable state. The first equation holds because **z** is a deterministic function of **x**.

Equation 1 is the foundation of our *probabilistic tokenization theory* for protein structures. It justifies that a discrete prior (denoting the metastable states) for continuous protein structures is reasonable. It also implies that, if the metastable states are few (given a large *τ*_obs_), **ϵ** should account for large and complicated variations within each state. On the contrary, if the metastable states are defined at a small timescale,**ϵ** is only responsible to very subtle structural variations within each state, but the number of **z** could quickly explode. Put it in other words, there is a trade-off between the *compactness* of **z** (i.e., the number of defined metastable states) and the *informativeness* of **z** (i.e., the residual intra-state variation that has to be explained by **ϵ**), as illustrated in Fig. 1a.

Inspired by the lattice model for proteins(Taketomi et al., 1975), it is plausible that the number of metastable states grow with the length of proteins, so we can assign a finite number of discrete states to each residue (namely, tokens), and the combination of these residue-wise tokens, define the overall state of the protein. From this perspective, amino acid(s) is indeed one kind of such token, i.e., *Anfinsen’s token*(s). Anfinsen’s tokens are *probabilistic* in nature because they do not correspond to a single snapshot of protein 3D structure, but to all the folded structures with *τ*_obs_ ≈ *τ*_fold_ according to the Anfinsen’s hypothesis. However, Anfinsen’s tokens are extremely compact (with a small vocabulary size of only 20), thus, leaving the intrastate variations large and hard to estimate. That explains why conformational prediction based on amino acids is a tremendously challenging task. Due to the absence of an effective *detokenization* algorithm (although significant advance has been made since AlphaFold(Abramson et al., 2024, p. 3; Jumper et al., 2021; Senior et al., 2020)) which can trustworthily map back amino acids to the folded conformations of the protein, they are often not regarded as a 3D representation for protein structures.

Compared to Anfinsen’s tokens, tokenizing metastable states at finer timescale *τ*_obs_ < *τ*_fold_ has several compelling advantages: i) more detailed changes in structures corresponding to functional switch can be described including alternative conformations; ii) tokens can strike good balance between being compact and being informative; iii) a more efficient detokenizer can be obtained to backmap the tokens. Specifically, given a protein 3D structure **x**, we tokenize the metastable structure ensemble **{x}** associated with **x** into probabilistic, amino acid-like tokens, which can be detokenized back to conformations from the corresponding metastable state.

#### 2. ProTokens: Aritificial amino-acid tokens can be extracted through unsupervised deep learning

Due to the fact that the relaxation of side chains is often much faster than the backbones, metastable conformations of the backbone and sidechain can be tokenized separately(Monticelli et al., 2008). The token for an all-atom protein 3D structure ensemble, termed as ProToken, is thus defined by the Cartesian product of the backbone token and the sidechain token.

A *wildcard backbone token* is introduced: When it is combined with any sidechain token in the form of amino acid, it becomes a valid ProToken equivalent to Anfinsen’s token. Particularly, this special subset of ProTokens degenerately encodes all the folded conformations of the protein. Therefore, ProToken unifies the modality of 1D and 3D representations of protein structures. In addition to the unified modality, we also equip each ProToken with *Janus representations*, that is, two mutually mappable representations: One is a discrete token index, which is amenable for language-based foundation models; The other is a continuous symmetry-free token embedding, which is ready for diffusion-based foundation models.

We devise an unsupervised deep learning approach called *ProToken Distiller* to extract ProTokens automatically from experimental and physics-simulated data. Inspired by the profound connections between ProTokens and amino acids, the *ProToken Distiller* is a joint optimizer for the classical inverse folding and the structure prediction problems (Fig. 1b), except that the conventional predefined amino acid vocabulary is replaced by a learnable set of ProTokens so that the gradient flow can be back-propagated end to end.

We train the ProToken Distiller using a curated protein structure dataset(Liu et al., 2022) consisting of cleaned single chains collected from the Protein Data Bank(Berman et al., 2000) with a truncation date of Oct. 13th. 2021 (prior to CASP14). We filter these structures and keep chains without unresolved structural gaps, remove structures shorter than 30 residues and those from NMR experiments (due to the possibility of lacking metastability), and hold out over 500 structures for validation and use the others (∼552 K) as the training data. During training, each single chain is cropped up to 256 residues.

We initialize the codebook of learnable backbone tokens with a size of *K* = 768. During training, the codebook size is reduced to *K* = 513 through variational clustering including a default wildcard token. The number of sidechain tokens is fixed to 20 without further optimization, thus, resulting in a total number of 513 × 20 = 10260 ProTokens.

The dimensions of the backbone and sidechain token embeddings after compression is 32 and 8, respectively, leading to a final dimension of 40 for each ProToken embedding. More detailed training settings can be found in SI Section II-A.

Because the sidechain parts in ProTokens are set as default and not deliberately optimized, to keep notations simple and consistent, in the following sections regarding the benchmark of optimized ProTokens, unless specified otherwise ProTokens are used interchangeably with backbone tokens, and protein structures refer to the backbone structures.

### B. Variationally optimized ProTokens are informative and generalizable representation of protein structures

#### 1. ProTokens preserve both global and local patterns of metastable protein 3D structures

To demonstrate the efficacy of ProTokens for 3D structure representation, especially for backbone conformations, we evaluate the quality of the reconstructed backbone protein structures decoded from the backbone ProTokens against the ground-truth structures in the test set. Provided that the Decoder is a generative model which may yield different structures given the same ProToken, during benchmark we turn off any randomness so that the model runs deterministically.

The test set contains 87 and 44 single chain targets from CASP14 and CASP15 which are publicly accessible. These test data are used to reasonably assess the quality of backbone ProTokens. According to Fig. 3a, the average TM-score of reconstruction (rTM-score) exceeds 0.96 and the median rTM-score exceeds 0.97 for both test sets. Note that such subtle deviations fall in the range comparable to the intrinsic fluctuations of a metastable protein structure and are considered as adversarial perturbations during the training of ProToken Distiller. In conclusion, this experiment demonstrates that ProTokens can sufficiently preserve the overall patterns of the protein backbone structure. Beises, we also provide examples of protein structures (Fig. 3d) which can be reasonably detokenized from ProTokens, but not from the amino acid sequences, confirming that ProTokens are more informative than Anfinsen’s tokens.

**Fig. 3.**
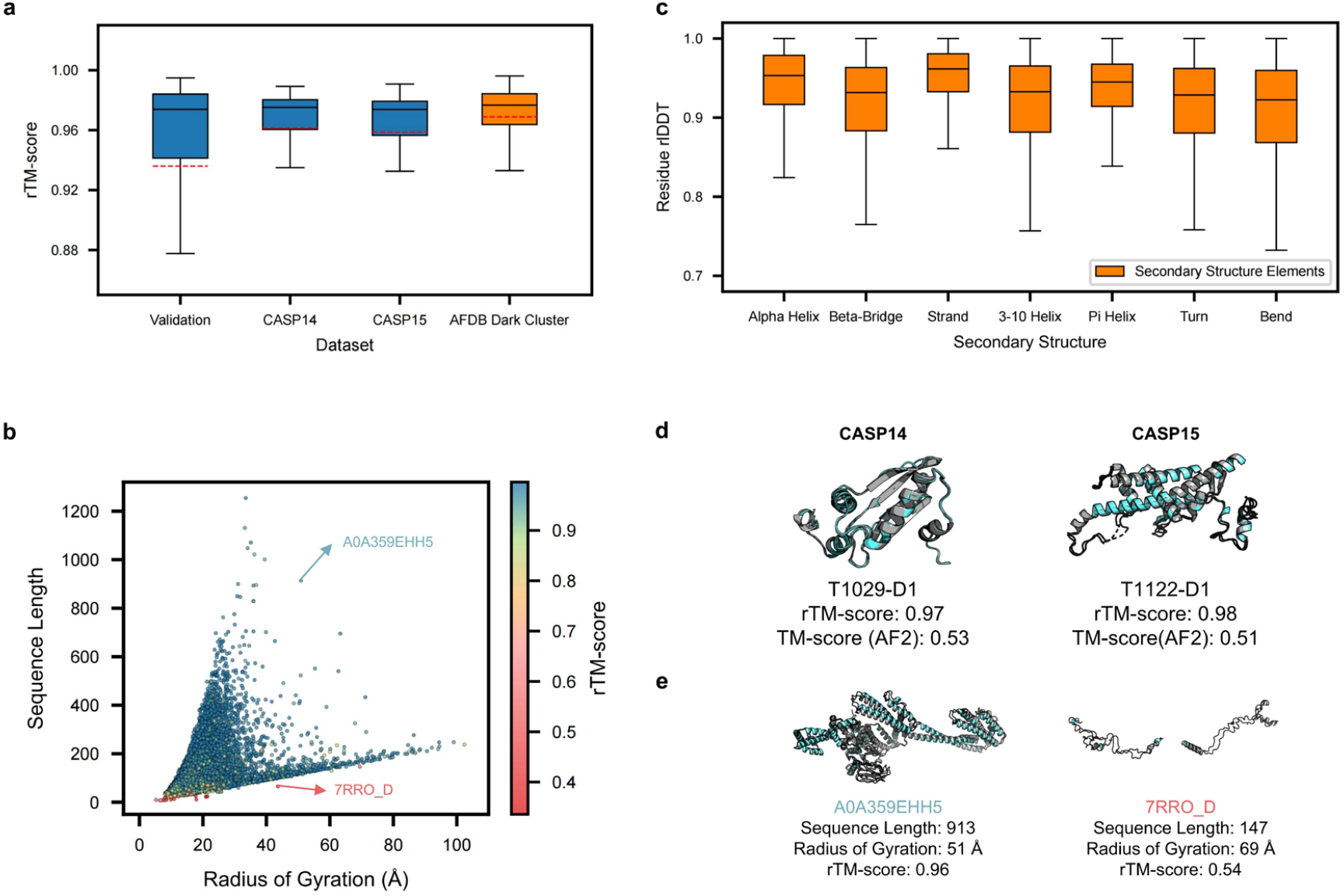
ProTokens are informative representations for tertiary structures. **a**, Statistics of the TM-score between the decoded structure from ProTokens and the reference structure, rTM-score, over the validation and test sets. **b**, ProTokens are relatively robust to the length and shape of proteins, but susceptible to extremely extended structures. **c**, ProTokens are locally accurate by characterizing various secondary structure elements well. **d**, Examples of ProToken-decoded structures (cyan) superimposed with the experimental reference (gray), which are more consistent than Anfinsen’s token-decoded structures measured in TM-score. **e**, Comparison of examples when ProTokens succeed (left) or fail (right) in describing the tertiary structures. The box plots in **a** and **c** are defined by the median as the centre black line, first and third quartiles as the box edges and 1.5 times the inter-quartile range as the whiskers. The red dash lines plot in **a** represent the mean value.

Next, we wonder whether the efficacy of ProTokens is sensitive to the length or shape of a protein, so we zoom in over the results of the test set in scatters (Fig. 3b). In general, the reconstruction quality is relatively consistent with respect to the length and shape (in terms of the radius of gyration) of the protein. However, when the protein exhibits a relatively extreme shape, for instance, very extended or in a nearly-linear form (Fig. 3b), the fidelity (defined as the reconstruction quality) of ProTokens drops significantly regardless of whether the protein is short or long. Considering that extremely extended and linear protein structures mostly contain a limited number of stabilizing interactions (Fig. 3e), such geometry usually lacks sufficient stability in physiological conditions. This phenomenon implies that the efficacy of ProTokens may be susceptible to the stability, but not to the length or shape, of a protein structure.

We further examine whether ProTokens discriminate against certain local structural patterns, particularly, those local elements that are less abundant in the training data. It can be concluded from Fig. 3c that ProTokens can reasonably reconstruct the local environment (in terms of residue local distance difference test, lDDT(Mariani et al., 2013)) of residues taking various second-order structures, showing no preference or discrimination over specific patterns.

Given the reliable reconstruction quality of ProTokens, we released a compressed PDB dataset (PT-PDB), containing ∼550k single chains with the backbones being compressed and stored in ProToken indices, leading to a compression ratio of 24. We also released the embeddings for the ProTokens, based on which the compressed backbone structures can be decoded back to Cartesian coordinates of atoms.

#### 2. ProTokens generalize to the dark protein universe, disconnected domain assemblies and multimers

The test set in the above experiment consists exclusively of structures from experiments, which may be subjected to human biases. Because ProTokens are trained solely on experimental structures, it is possible that such biases are inherited. Based on an AI-predicted protein structure dataset (AFDB)(Varadi et al., 2024), previous research reveals that there is a large volume of unexplored dark space of the protein universe(van Kempen et al., 2024). Therefore, we extract high-confidence structures from the AFDB dark cluster dataset and interrogate ProTokens’ capability of reconstructing these out-of-distribution (OOD) folds. Fig. 3a shows that these OOD single-chain structures can be represented by ProTokens as well, and the reconstruction quality shows no significant difference from that of the experimental targets statistically. This experiment demonstrates that ProTokens are generalizable for tertiary structural patterns.

Next, we wonder whether ProTokens are able to capture quaternary structural patterns, despite the fact that no such examples were presented during training. Considering that multi-domain proteins are often regarded as transitioning from tertiary to quaternary structures, we first collect several samples containing multi-domain chains from CASP14 and CASP15 test sets. These single chains are manually chopped into annotated domains according to the official definition, yielding a set of disconnected multi-domain assemblies which resemble protein complexes. Noteworthy, the training set of ProTokens merely contains single-chain structures without discontinuity (i.e., without structural gaps), so these multi-domain assemblies are also OOD examples. Nevertheless, we allow ProTokens to treat them by manually setting a residue index gap between domains during inference. We surprisingly find that ProTokens can reconstruct the assembled structures reasonably well (Fig. 4a) when the inter-domain contacts are not sparse.

**Fig. 4.**
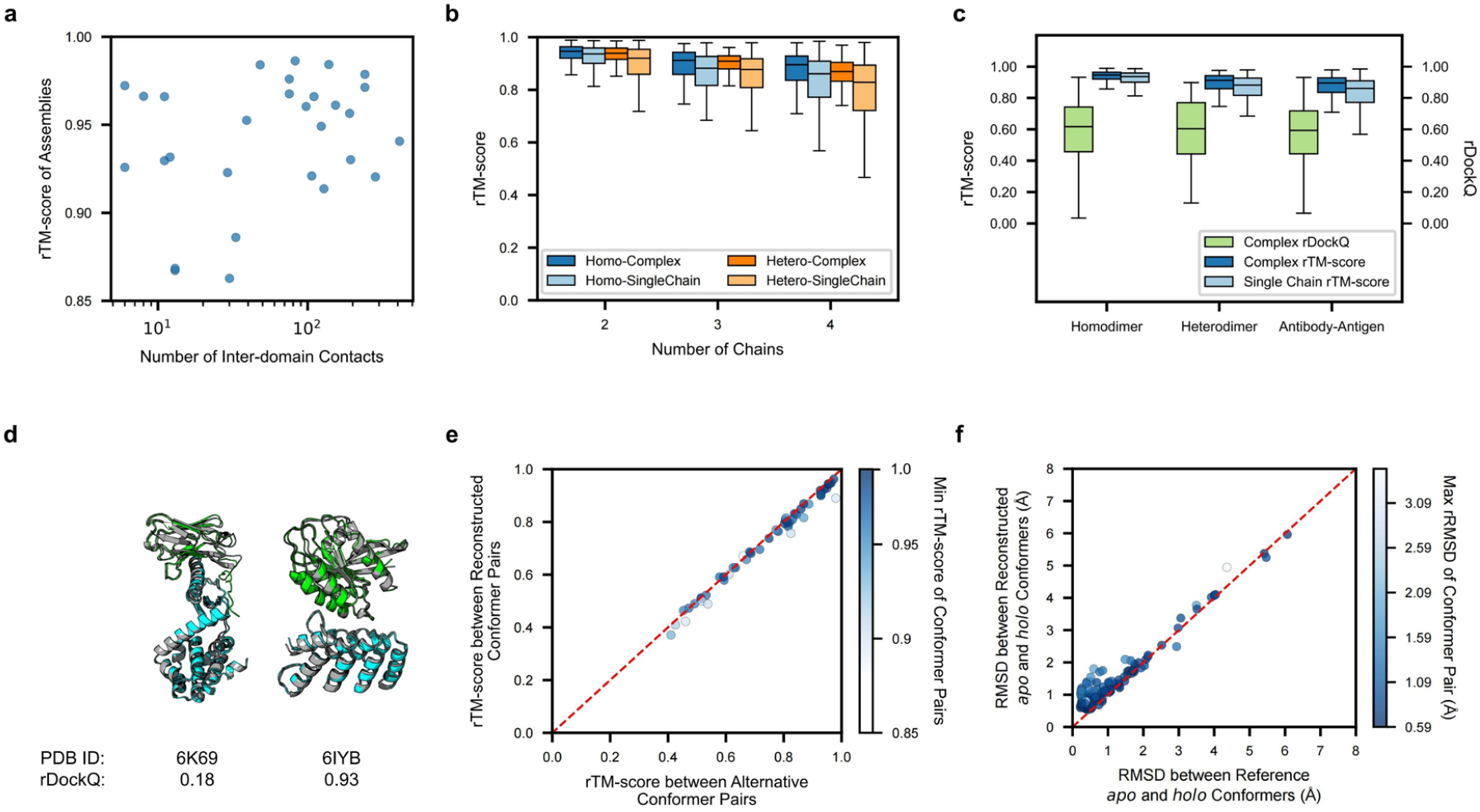
ProTokens generalize well to quaternary structural patterns. **a**, ProTokens can reconstruct in high quality the disconnected domain assemblies with sufficient inter-domain interactions. The sample size is 28 in total. **b**, ProTokens characterize both the tertiary and quaternary structure patterns in protein complexes. **c**, The interface between homo- and hetero-dimers can be reasonably reproduced by ProToken. **d**, Compared to heterodimer (PDB ID: 6IYB), degraded fidelity is observed for the antigen-antibody interface (PDB ID: 6K69) with sparser quaternary interactions. The aligned structures are shown with the groundtruth colored in gray, domain 1 in green, and domain 2 in cyan. **e**, Assessing the fidelity of ProTokens for representing alternative metastable conformations. **f**, Assessing ProTokens’ capability of distinguishing *apo* and *holo* conformers for ligand-binding proteins. The box plots in **b** and **c** are defined by the median as the centre black line, first and third quartiles as the box edges and 1.5 times the inter-quartile range as the whiskers

Based on this encouraging result, we further test ProTokens over real-world protein complexes, including homomers and heteromers. The overall performance is satisfying in that both the single-chain and the complex structure can be well reconstructed (Fig. 4b). This result confirms ProTokens’ capability of characterizing both the tertiary and quaternary structure patterns, although slight performance degradation is observed when the protein complex involves more chains (Fig. 4b).

In addition to the overall shape, the accuracy of the complex interface is of particular interest when assessing a multimer structure. Motivated by this, we compute the DockQ score(Basu and Wallner, 2016) for the tested dimers, including antigen-antibody (Ag-Ab) complexes(Gao et al., 2022). Figure 4c shows that the interface is relatively accurate(Basu and Wallner, 2016), though not perfect, with an average and median DockQ exceeding 0.49. Noteworthy, compared to multi-domain assemblies or heterodimers, we find that the reconstructed Ag-Ab complex interface shows larger variance, although each single chain is reconstructed well (Fig. 4d). This may be due to the fact that the interaction patterns between Ag-Ab are often sparse and largely determined by the side chains, hence, are less comparable to the tertiary structural patterns that ProTokens are trained on. Besides, the flexibility of the Ag-Ab interface may also impact the fidelity of ProTokens.

Nevertheless, these experiments on multi-chain structures validate the generalization of ProTokens, which are trained solely on tertiary structures, over quaternary interactions. Our observation also implies that the quaternary structure patterns between protein chains may share deep connections with tertiary patterns within a chain, both arising from fundamental physics interactions between amino acid residues.

#### 3. ProTokens are descriptive of finer functional protein conformations

Researchers are particularly concerned with alternative conformations that may lead to functional switching of a protein. Despite many efforts being paid(del Alamo et al., 2022; Kalakoti and Wallner, 2024; Wayment-Steele et al., 2024; Zhang et al., 2023b), decoding alternative protein conformations according to the amino acid sequence remains an unresolved challenge. According to the probabilistic tokenization theory, such difficulty is largely due to the compactness of Anfinsen’s tokens (Fig. 1a), leading to large and multimodal intra-state variation (i.e., *p*(**ϵ**|z) in Eq.1) which is hard to model. Compared to the Anfinsen’s tokens, ProTokens are designed to be more informative by compromising its compactness. In principle, different metastable conformational ensembles of one protein, which are degenerate in terms of Anfinsen’s tokens, can now be distinguished by ProTokens.

We thus conduct experiments to test whether ProTokens are able to characterize alternative conformations of a protein. We first prepare a test set consisting of proteins with a pair of alternative conformations from 50 PDBFlex(Hrabe et al., 2016) clusters with the local RMSD larger than 2.0 Å. Note that these conformers are all resolved by experiments, showing sufficient stability. We then calculate the reconstruction quality for each pair of the alternative conformers and find that ProTokens can not only characterize well the structure of each conformer, but also authentically preserve the relative difference between a pair of conformers (Fig. 4e). This experiment verifies the capability of ProToken as informative representations for finer-scale conformations of a protein.

Many proteins are known to undergo conformational changes or adaptations during binding to ligand(s), we thus wonder whether ProTokens can characterize the binding-altered backbone conformations of a protein. Similar to the previous experiment, we test the reconstruction quality of a set of proteins with different *apo-* and *holo-*form conformations (defined as conformers mutually different with a minimum backbone RMSD larger than 2.0 Å with and without ligand binding). According to the results shown in Fig. 4f, ProTokens are able to faithfully describe both the *apo* and *holo* conformers of a ligand-binding protein and preserve their relative differences. This important feature permits the use of ProTokens for the design of ligand-binding proteins involving conformational changes.

### C. Probabilistic tokenization yields robust and interpretable encodings of metastable protein backbone structures

#### 1. Similarity in ProTokens parallels similarity in 3D structures

It is compelling to analyze the reasons behind ProTokens’ ability in representing protein 3D structures. A reasonable hypothesis is that the structure similarity may be reflected by ProTokens, and we start our analysis from the ProToken embeddings. Provided that each residue in a protein 3D structure is assigned with a ProToken, we wonder whether the ProToken embedding of the residue characterizes its local structure environment, inspired by the classical word embedding theory(Mikolov et al., 2013).

On the one hand, we compute the cosine similarity between each pair of ProToken embeddings (Fig. 5a), which reflects the token similarity learned and determined by the optimized model. On the other hand, we compute the BLOSUM (short for blocks substitution matrix)(Henikoff and Henikoff, 1992) of ProTokens using residue pairs with similar local environments by convention. These residue pairs are drawn from a dataset specifically for developing structure alignment methods like FoldSeek(van Kempen et al., 2024). A higher BLOSUM coefficient between two tokens indicates a higher chance of them being shared by two residues with similar local structural environments. Intriguingly, we find that the BLOSUM of ProTokens correlates significantly well with the similarity matrix of ProToken embeddings (Fig. 5a). This result reveals an important feature of ProToken: if two residues adopt ProTokens with high similarity, they are likely to be positioned in a similar 3D structural environment.

**Fig. 5.**
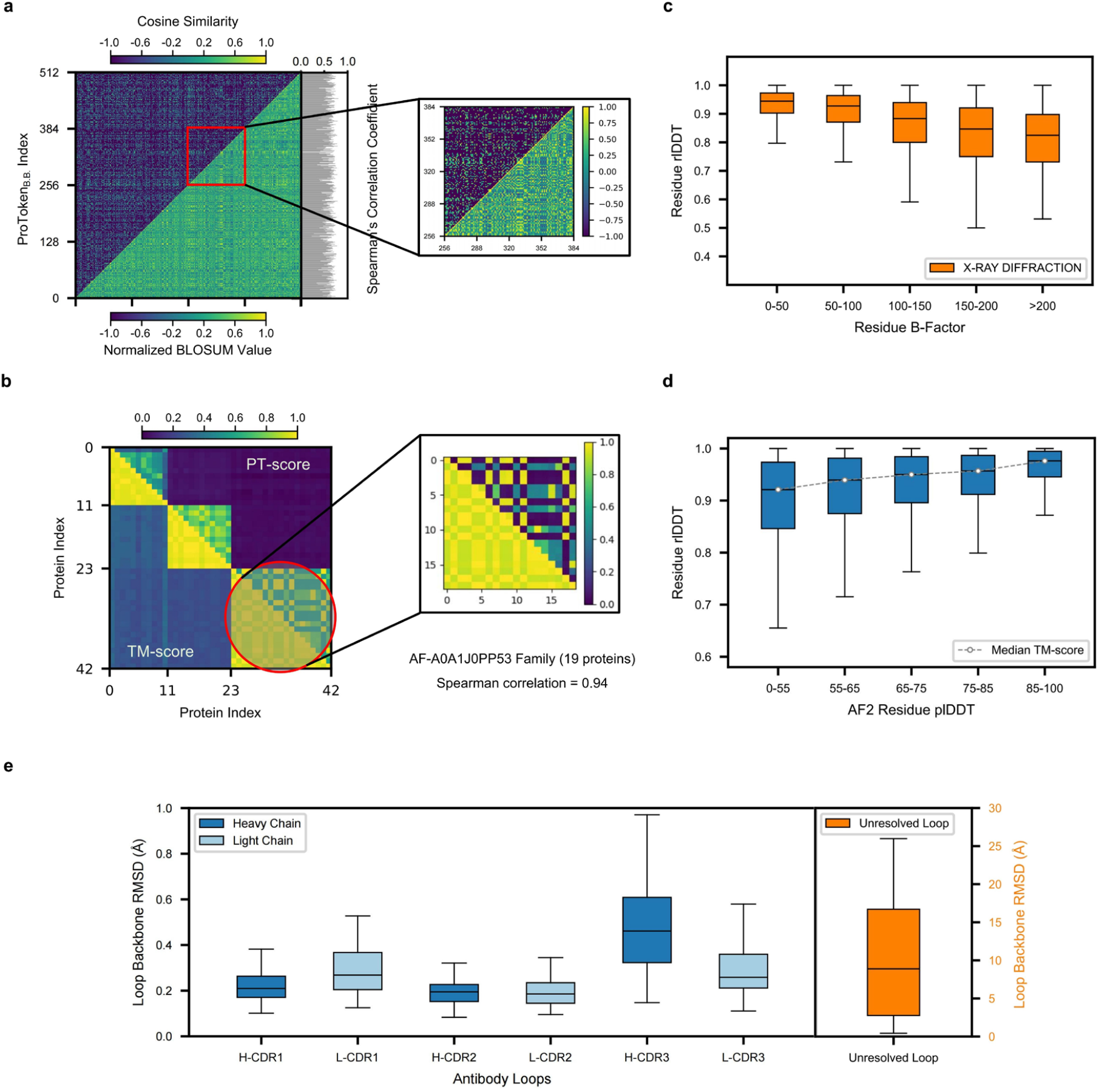
ProTokens reflect similarity and metastability of protein structures. **a**, The cosine similarity matrix between ProToken (upper left) embeddings correlates significantly with the BLOSUM (lower right). **b**, PT-score (defined as the difference in ProTokens) correlates quantitatively with TM-score, and can be used for structure comparison and clustering. **c**, The fidelity of ProTokens correlates negatively with the flexibility of local structures as measured by experimental B factors. **d**, The fidelity of ProTokens correlates positively with the stability of local structures as indicated by AlphaFold2 plDDT values. **e**, Antigen-binding conformations of antibody loops are represented well by ProTokens (left panel), indicating metastability, in contrast to experimentally unresolved loops (right panel; note the scale of y-axis is different from the left panel).

Since the residue-wise similarity in ProTokens has been proved to characterize local structural similarity, it is likely that the sequence-wise distance measured by ProTokens may also quantify the overall structural difference. We thus collect a set of clustered structures AFDB(van Kempen et al., 2024; Varadi et al., 2024), calculate the TM-score as well as the ProToken Similarity Score (PT-score) between each pair of the structures. It can be concluded from Fig. 5b that a higher PT-score between proteins indicates a higher structural similarity measured by TM-score. Specifically, PT-score is not only able to distinguish similar structure pairs from unsimilar ones, but also display a quantitative correlation with TM-score so that it can reasonably sort and rank structures according to the similarity with respect to a reference structure. Therefore, the PT-score can be a promising alternative option of quantifying the structural difference between proteins that is required by structure search and clustering algorithms.

#### 2. Fidelity of ProTokens reflects the (meta)stability of 3D structures

As observed in the previous section, the fidelity of ProTokens seem to correlate with the stability of a protein structure (Fig. 3b), possibly due to the probabilistic tokenization process, that is, the metastable structure patterns are prioritized by ProTokens over the transient and dynamic snapshots of a protein 3D structure. To further investigate this phenomenon, we zoom in our analysis over protein residues, considering that even within a single protein, the stability of different residues may be heterogeneity. For instance, it is common that some substructures are intrinsically more dynamic or less stable than the other regions, which may result in less confident or even unresolved parts in experimentally determined structures.

Therefore, we disassemble the test samples into residues, and categorize them according to the B-factors reported by the crystallization experiments. In general, a larger B factor indicates larger intrinsic dynamics (or less stability) of the residue in its 3D structure(Sun et al., 2019). Not surprisingly, we find that the fidelity of a ProToken corresponding to a certain residue correlates with the B-factor of that residue (Fig. 5c). In other words, if the local structure is less stable, it is also more likely to be smeared by ProTokens. We further corroborate this conclusion based on another test set consisting of AF2-predicted structures. Previous research revealed that the predicted confidence (predicted lDDT) of AF2 can reflect to a certain degree the structural flexibility of the residue(Ma et al., 2023). In line with previous findings, we observe that on average, when the plDDT of a residue lowers, the fidelity of ProTokens for that residue gets worse (Fig. 5d).

Additionally, as many loop or intrinsic disordered regions of the protein usually cannot be resolved by crystallization due to lack of metastability, it is likely that ProTokens are not able to represent them either. To test this hypothesis, we manually fix and fill the unresolved regions of the experimentally determined structures using PDB-Fixer(“openmm/pdbfixer,” 2024) and AF2, then compute the reconstruction quality of ProTokens for these regions. Consistent with the previous conclusion, we observe a significant drop in the fidelity of ProTokens for these disordered regions (Fig. 5e). However, we notice that, unlike the manually fixed random regions, the experimentally resolved loop conformation of an antigen-bound antibody can be well reconstructed from ProTokens (Fig. 5e). This observation implies that the antigen-binding loop of an antibody is a metastable conformer of a free-form antibody, in line with the thermodynamics theory of protein binding. Unlike intrinsically disordered loops, the difficulty in experimentally resolving these metastable loop conformers alone may result from the existence of other competing metastable conformations.

## 3. Discussion

Proteins serve as the most fundamental and important functional molecules for life. Understanding the relationship between the constituent of a given protein and its function remains central in various research scopes. It is commonly assumed that the function of a protein is largely determined by its 3D structure, and that the structure is mostly dictated by its amino acid sequence. Protein structure thus plays a central role linking its sequence and function, and two widely concerned problems naturally arise: 1) How to predict the functional protein structure given its sequence, and 2) how to generate proteins (including structures and sequences) to fulfill a specific function. AlphaFold has achieved remarkable progress on the former challenge, however, the latter question remains wide open and is still calling interest from an active research community for protein design.

In principle, protein design requires joint generation of a protein backbone structure and side-chains which are mutually compatible, that is, the sequence can well fold into the designed structure. However, in common practice, the generation process is factorized: a protein backbone structure is first designed according to the desired functions, then an amino-acid sequence is assigned in order to stabilize the designed structure. Such factorization rests on a strong but often overlooked assumption of “designability”, namely, there exists at least one sequence combo of (natural) amino acids that can fold dominantly if not exclusively to the designed protein structure. However, although structures can be generated with arbitrary variety in the coordinate space, these backbones may not be stabilized by any combination of natural amino acids, resulting in failure of the design. Unfortunately, the high failure risk of many factorized design approaches reflects the fact that our understanding of the “designable space” of protein structures is still quite limited. Therefore, proper modeling of the designable space of protein structures is urgently needed to improve the success of factorized protein design.

Recently, data-driven generative methods are gaining increasing attention and provide new opportunities to this long-standing challenge. However, 3D protein structures are composed of atomic coordinates which exhibit specific symmetries and are subjected to physics priors, such as transrotational equivariance and polymer restraints, hence, significantly impeding the transaction of modern generative AI for protein structure design. In a quite different approach, protein can also be designed by directly generating the amino acid tokens. In contrast to the 3D structure, the amino acid sequence, also known as the 1D structure of a protein, is formally discrete and SE(3)-invariant. Therefore, the amino acid sequence becomes a compelling candidate as input to language models. Many methods have been developed for protein understanding and designing tasks by educating the LLMs over amino acid sequences. Unfortunately, the modality difference of protein structure representations (1D v.s. 3D) causes significant divergence in the research paradigms of protein design. Although both paradigms have witnessed a lot of advances, little has been investigated about the connections between the continuous and discrete representations of protein structures, and whether they can be interpreted by a unified theory, hence, eliminating the practical differences in 1D- and 3D-based protein design.

In this study, we proposed probabilistic tokenization theory and provided a unified perspective for the 1D- and 3D-based protein structure representations. The probabilistic tokenization theory underpins the feasibility of transforming continuous metastable protein structures into discrete tokens. Particularly, amino acids are special Anfinsen’s tokens corresponding to long observation timescale. Furthermore, we developed an unsupervised and variational optimization strategy, which integrates the structure prediction and the inverse folding tasks, to encode protein structures into an expanded vocabulary in addition to amino acids, called *ProTokens*, and detokenize them back to 3D coordinates. With the implementation of several important regularizers based on information theory, the optimization process ensures the sufficiency, necessity as well as robustness of ProTokens as representations of metastable protein structures.

We showed that discretizing protein structures with proper perplexity can lead to informative and relatively compact ProTokens. Using limited vocabulary of tokens, ProTokens can reconstruct 3D coordinates with high fidelity and reduce the trans-rotational equivariance as well as the polymer restraints of protein structures. We also find that ProTokens trained on single-chain dataset can generalize to multi-chain complex structures, indicating the transferability of basic physics interaction patterns that govern the tertiary and quaternary protein structures. More intriguingly, although ProTokens are able to describe alternative conformations (particularly function-related conformations) of proteins, including ligand binding sites and antigen-bound antibody loops, we find ProTokens cannot reproduce well the experimentally unresolved or ultra-dynamic regions of a protein such as those with high B-factors. These findings demonstrate that by means of probabilistic tokenization, ProTokens are good at describing metastable conformations that exhibit a relatively high stability, rather than overfit to transient structure snapshots that lack stability. Therefore, it would be an appealing direction to further investigate how the detokenized fidelity of ProToken correlates with the foldability of a protein backbone structure in the future.

Last but not least, we speculated on possible pitfalls for the usage of a transformed protein structure representation such as ProTokens for downstream tasks. The robustness of the transformed tokens is vital to understanding tasks like function prediction because subtle fluctuations of protein structures mostly do not change its function. On the other hand, the degeneracy or duplicate tokens could be very harmful to the latent generative model based on ProTokens due to a poor MLE bound. We also introduced guidelines to ameliorate these issues. Based on these foundations, we expect that ProTokens will enble transaction of modern generative AI for protein annotation and protein design in the near future.

## 4. Methods

### A. Pitfalls of a Machine-Learned Protein Structural Representation

After transforming the spatial coordinates of metastable structures into SE(3)-invariant (continuous or discrete) representations like ProTokens, one may apply them for downstream tasks related to protein structures. However, we reveal that high risk exists if the transformed representations are not optimized or implemented with caveats, and summarize several potential pitfalls of learning and applying a transformed representation. Awareness of these issues are reflected in the specifically designed model components and training objectives which will be elaborated in the following sections.

#### 1. The space of the transformed representation should be compact

Taking ProTokens of a *N*_res_-long protein as example, the real-valued token embeddings take the shape of (*N*_res_, *d*). If 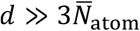 (where 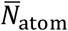 stands for the average number of atoms per residue), diffusion in the transformed embedding space will be less efficient (both in terms of data and training) than spatial diffusion approaches like RFDiffusion and AlphaFold3(Abramson et al., 2024; Watson et al., 2023), provided that the likelihood of the model decays exponentially with the dimensionality of the representation.

Besides, the number of all possible combinations of ProTokens grows combinatorially with respect to the vocabulary size *K*. Since the number of foldable protein structures (as well as functionally relevant metastable states) is unlikely to exceed 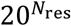 the reasonable number of tokens used during training should lie in the range between tens to a few hundreds.

To extract compact ProTokens, we design a *Compressing* module (see Section II-C and SI Section I-C. for more details) in the *ProToken Distiller* to properly compress the ProToken embeddings lengthwise or depth-wise. Besides, we derive a variational information bottleneck (VIB)(Alemi et al., 2019) to quantify the necessity of ProTokens, and the vocabulary size *K* is optimized with respect to the VIB loss by *variational clustering* during training (Eq. 11; see Section II-D and SI Section I-C for more details).

#### 2. The transformed representation should be robust against intrinsic structure fluctuations

In terms of protein physics, fluctuations are intrinsic to metastable protein structures, which may cause subtle structure changes but do not alter the function. One particular concern of a learned structural representation is that the *robustness* of the yielded embeddings and tokens against subtle structure perturbation may not be guaranteed (Fig. 1c), while the susceptibility to intrinsic fluctuations will be harmful to downstream tasks such as function predictions.

To alleviate this issue, we introduce adversarial examples by adding physics-informed fluctuations to any input structure, and adversarially train the model to behave robustly to the *adversarial attacks*. Besides, we also train a *Deduplicator* module to relax fluctuated structures within a metastable ensemble towards a single stable representative structure, and collapse the embeddings of the fluctuated structures (see Section II-C and SI Section I-B for more details).

### 3. Existence v.s. uniqueness: The degeneracy of transformed representation should be addressed

Noteworthy, the *uniqueness* of a learned ProToken corresponding to a certain structure x cannot be guaranteed through data-driven training. *Duplicate* or degenerate ProTokens may exist that can be decoded to (almost) the same structure (Fig. 1c). Degeneracy is particularly poisonous when ProTokens are used for maximum likelihood estimation (MLE) of protein structures by generative models such as auto-regressive and diffusion models. To be more specific, after transforming the representation of protein structure **x** to ProToken **z**, which can be detokenized back via a decoder **x** = *g*_*ϕ*_(**z**), the (log-)likelihood of the structure **x** should be computed in the following way as in latent generative models(Rombach et al., 2022; Yu et al., 2022),

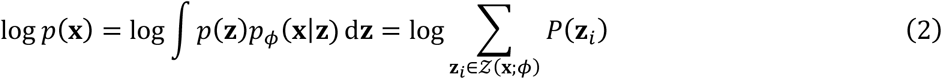

where *p*_*ϕ*_(**x**|**z**) = *δ*(*g*_*ϕ*_(**z**), **x**) is defined as “*duplicate distribution*”, which consists of all ProTokens **{z}** that can be decoded back to **x** through the decoder. Since **z** is discrete, integration of *p*_*ϕ*_(**x**|***z***)*p*(**z**) over **z** is equivalent to the summation of *p*(**z**_*i*_) over a finite and countable *duplicate set* 𝒵(**x**; *ϕ*).

Since we barely have information about 𝒵(**x**; *ϕ*) *a priori*, the exact likelihood in Eq. 2 is hard to compute in practice. To address this issue, we derive computationally efficient lower bounds (Eq. S4 and Eq. S7) for the likelihood and design a *Duplicator* module (see Section II-C for more details) to explore and expand the duplicate set to reduce the bias and variance for likelihood estimation.

### B. Models

Tokenizing metastable states associated with a given protein structure can be cast as a *conditional generative learning problem* as in invertible coarse graining(Zhang et al., 2023a). We design a deep neural network system, *ProToken Distiller*, to achieve this goal (Fig. 2a).

#### 1. Probabilistic conditional decoder

Specifically, we aim at constructing a metastable conformational distribution *p*_*ϕ*_(**x**) and sampling from it according to a structure **x**, which can be achieved through a probabilistic conditional *Decoder g*_*ϕ*_ (Fig. 2b) using the reparameterization trick as in VAE(Kingma and Welling, 2022) or GAN(Goodfellow et al., 2014),

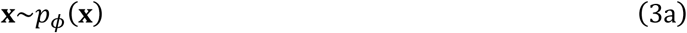

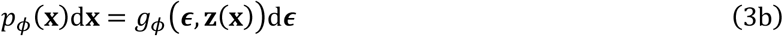

where **ϵ** is random noise from a known prior like Gaussian distribution, and **z(x)** is the embedding of a tokenized **x** provided as the conditional information to the *Decoder*. According to the probabilistic tokenization theory (Eq. 1), the ProToken **z** specifies the identity of the metastable state, whereas **ϵ** accounts for the conformational fluctuations within the state.

The Decoder is a composite function consisting of a *Token Duplicator* module and a *Detokenizer* module: The *Token Duplicator* is responsible to expand and sample from the duplicate set in Eq. 2, whereas the *Detokenizer* module is a SE(3)-equivariant generative model which samples protein structures from a metastable ensemble corresponding to a given ProToken string. More details about the *Token Duplicator* module and *Detokenizer* module can be found in SI Section I-A.

The conditional embedding **z(x)** = *h*_*θ*_ *ο f*_*θ*_(**x**) is obtained via a composite of *Encoder f*_*θ*_ and *Tokenizer h*_*θ*_, which transforms the all-atom protein 3D structure **x** into SE3-invariant embeddings of discrete tokens.

#### 2. SE(3)-invariant encoder

The *Encoder f*_*θ*_ comprises an SE(3)-invariant *Structure Encoder* module (Supplementary Fig. 3), and a *Deduplicator* module (Fig. 2c). As explained, the *Deduplicator* module is introduced to improve the *robustness* of the yielded embeddings against intrinsic structure fluctuations (see SI Section I-A for more details about the *Duplicator*). Given the separation of timescales of backbone and sidechain motions, the sidechain and backbone structures are encoded separately,

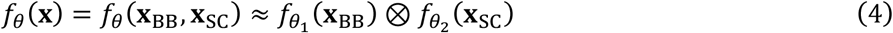

where **x**_BB_, **x**_*sc*_ denote the backbone and sidechain structures, respectively; and ⊗ denotes Cartesian product (i.e., concatenation of tensors). The resulting *f*_*θ*_(**x**) is a continuous SE(3)-invariant embedding for the protein structure. More details about the backbone and sidechain structure encoders can be found in SI Section I-B.

Considering that metastable structure ensembles can be reasonably represented by discrete tokens, we prepend a Tokenizer *h*_*θ*_ to the Encoder in order to *variationally cluster* (or discretize) *f*_*θ*_(**x**) into quantized ProTokens.

#### 3. Variational tokenizer

The Tokenizer *h*_*θ*_ = *r*_*θ*_ *ο s*_*θ*_ discretizes the structural embeddings *f*_*θ*_(**x**) into ProTokens **z(x)** (Fig. 2d), consisting of a composition of a *Clustering* module *s*_*θ*_ and a *Compressing* module *r*_*θ*_ (see more details about the Compressing module in SI Section I-C). The *Clustering* module aggregates the continuous embedding learned by the Encoder into *K* clusters (i.e., “codes” or “tokens”), and each cluster is assigned with a *d*-dimensional vector as the cluster center (or token embedding). Since the backbone and sidechain embeddings are obtained through two independent tracks, the Clustering module also operates separately for backbone and sidechain embeddings,

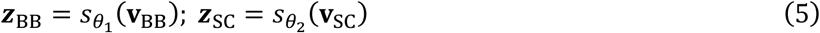

**z**_BB_, **z**_*sc*_ denote the backbone and sidechain tokens, respectively. The ProToken for all-atom structure is then assembled by Cartesian product **z** = **z**_BB_ ⊗ **z**_sc_. Each ProToken has *dual representations* which are mutually mappable: one corresponds to the discrete cluster index, the other is the embedding of the cluster center which lies in a continuous vector space.

To make the clustering procedure end-to-end differentiable, we approximate the gradient flow with straight-through estimators(Bengio et al., 2013; Liu et al., 2023) for backbone tokenization, which will be elaborated in the next section (Section II-D). Details about the sidechain tokenization can be found in SI Section I-C.

Noteworthy, there is a fundamental difference between the Clustering module and common VQ models such as VQ-VAE(Oord et al., 2018) or VQ-GAN(Esser et al., 2021): The clustering is performed *variationally*, that is, the number of alive codes should be as small as possible in order to tighten the variational information bottleneck (see more details in Section II-D), which contrasts sharply to state-of-the-art VQ training where a high usage of codes is usually preferred(Sun et al., 2024; Tian et al., 2024).

### C. Optimization of ProToken Distiller

Overall, the ProToken Distiller (*g*_*ϕ*_, and *f*_*θ*_ *ο h*_*θ*_) is optimized towards the following coupled objectives:

1. minimizing the divergence between *p*_*ϕ*_(**x**) and *p*_*D*_(**x**);
2. maximizing the mutual information between ProTokens **z** and *g*_*ϕ*_(**ϵ**; **z**(**x**)), while minimizing the mutual information between ProTokens **z** and the input structure **x**;
3. minimizing the divergence between *f*_*θ*_(**x**) and the encoding of the adversarial example 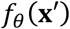

Intuitively, the first objective ensures the *sufficiency* of ProTokens as a transformed representation of metastable protein structures. The second objective guarantees the *necessity* and *non-redundancy* of the transformed representation. The last objective improves the robustness of ProTokens against intrinsic structural fluctuations. Technically, these objectives can be achieved by minimizing an InfoGAN loss(Chen et al., 2016) regularized by variational information bottleneck(Alemi et al., 2019) and adversarial training(Goodfellow et al., 2015). As a result, the final loss function *L*_PD_ for the ProToken Distiller to be minimized is a linear combination of the sufficiency loss *L*_suf_, necessity loss *L*_nec_, and robustness loss *L*_rob_ with a reasonable set of hyperparameters,

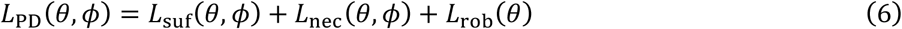

#### 1. Data preparation

In order to train the probabilistic Decoder, given each structure sample **x**_*D*_ from the training set, data augmentation is performed by means of metastable perturbation sampling (see SI Section II-A for more details), yielding a structure ensemble {x; **x**_*D*_} representing conformers from the same metastable state associated with **x**_*D*_. Samples from {**x**; **x**_*D*_} are used to compute the generative loss defined in *L*_suf_.

Furthermore, we construct adversarial examples 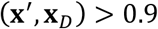 against **x**_*D*_ by setting a similarity cutoff TM-score 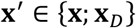 These examples are provided to the model for the calculation of *L*_rob_.

We note that both metastable conformers and adversarial examples can be prepared offline prior to the training, thus, incurring no extra overhead for training. Particularly, after the training proceeds, the adversarial examples can also be constructed online where the Decoder itself can serve as a perturbative sampler of an input **x**_*D*_. We will show that using these decoded structures as adversarial examples is indeed equivalent to the mutual information loss in *L*_nec_.

#### 3. Sufficiency

For the first objective, we adopt a loss function inspired by conditional GAN(Mirza and Osindero, 2014), which guides the *Decoder* to generate structures from the metastable states associate to an input structure **x**_*D*_,

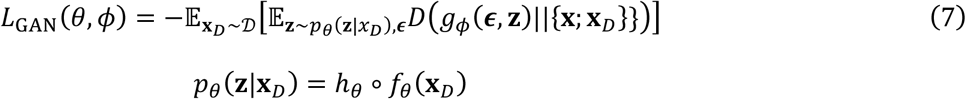

swhere *D*(⋅) is the critic measuring the divergence between two distributions; **z** is the token embedding (i.e., the identity of the metastable state) for **x**_*D*_ yielded by the Encoder and Tokenizer, and *p*_*θ*_(**z**|**x**_*D*_) depends on the code search algorithm during variational clustering (we adopt the nearest-code search(Oord et al., 2018) by default). **ϵ** represents random noises which are used to model the intrinsic structural fluctuations within the metastable state.

We repurpose the structure module of AF2 with the dropout trick(Wayment-Steele et al., 2024) for *g*_*ϕ*_(**ϵ, z**), where **ϵ** denotes the random dropout mask, allowing the SE(3)-equivariant structure module of AF2 to generate different structures conditioned on the same token **z**. Noteworthy, other unconditional generative models for protein backbone structures(Watson et al., 2023) can also be adopted and conditionally fine-tuned.

To optimize this GAN objective, we implement maximum-mean discrepancy (MMD) as the critic *D*(⋅)(Li et al., 2017), a non-parametric integral divergence metric between two distributions, which is known to stabilize and simplify the training of GANs(Li et al., 2017). The similarity kernel required by MMD is defined via frame aligned point error (FAPE)(Jumper et al., 2021), a Fréchet-like distance metric(Jing et al., 2024) for protein 3D structures.

In order to differentiate through the Clustering module *h*_*θ*_ in Tokenizer, we approximate the gradient flow of backbone tokens via the straight-through (ST) estimator(Bengio et al., 2013; Liu et al., 2023), and implement a commitment loss to reduce the errors of ST estimator,

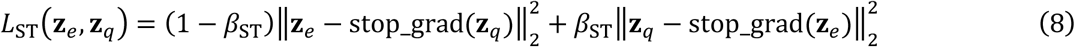

where **z**_*e*_, **z**_*q*_ represent vectors before and after quantization respectively. The final sufficiency loss for ProToken Distiller takes the following form,

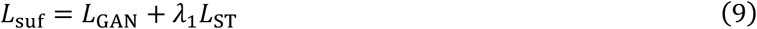

#### 3. Necessity

We perform *variational clustering* in the Tokenizer according to a VIB objective. Specifically, the training objective of *ProToken Distiller* can be recast in terms of VIB theory(Alemi et al., 2019),

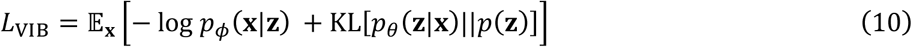

In VIB, *p*_*ϕ*_(**x**|**z**) is known as the prediction or reconstruction model (i.e., the Decoder), whereas *p*_*θ*_(**z**|**x**) is the inference model (i.e., the Encoder). By appending a Tokenizer to the Encoder, the prior *p*(**z**) and posterior *p*_*θ*_(**z**|**x**) become discrete, the VIB in Eq. 10 simplifies to,

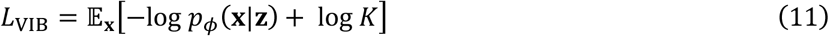

where *K* is the vocabulary size. Therefore, to minimize *L*_VIB_ is equivalent to gradually pruning the backbone token vocabulary and minimizing *K*, which is achieved by re-clustering the embeddings into a smaller number of clusters during training(Łańcucki et al., 2020).

Furthermore, to prevent the ignorance of conditional information (i.e., the ProTokens) by the generative *Decoder*, we also include a mutual information regularizer similar to InfoGAN(Chen et al., 2016), except that the auxiliary posterior estimator in InfoGAN is replaced by the conditional Encoder. This adaptation leads to a self-consistency term in the loss function,

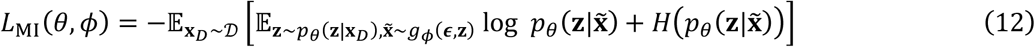

where the entropy term 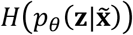 is similar to the entropy regularization introduced in VQ training(Chang et al., 2022). The final necessity loss for ProToken Distiller takes the following form,

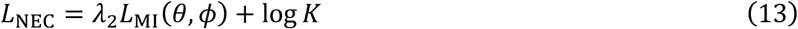

#### 4. Robustness

To help ProToken Distiller be better immune to adversarial attacks (i.e., structural fluctuations within a metastable state), we reuse the perturbative sampling data and present them as adversarial samples x′ ∈{**x**;**x**,_*D*,_} to the Encoder for each **x**_*D*_, and apply an adversarial training loss for the Encoder,

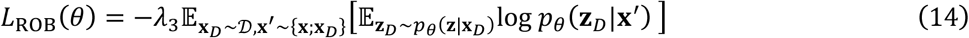

## Acknowledgments

This work was supported by the National Key R&D Program of China (No. 2022ZD0115002). The authors thank Dr. Sirui Liu and Dr. Xing Che for useful discussion, and gratefully acknowledge the support from Huawei MindSpore team to this research.

## Additional Information

### Conflict of Interest

Changping Laboratory is in the process of applying for provisional patent (Beijing Changping Laboratory) covering the discretization of biopolymer (including protein) 3D structures for the purpose of compressing, searching and generating 3D structures, that lists X.L., Z.C., J.Z. and Y.Q.G. as inventors. The other authors declare no conflict of interest.

### Code & Data Availability

Training set of ProToken is a cleaned version of the PSP Dataset which can be accessed via {http://ftp.cbi.pku.edu.cn/psp/true_structure_dataset/pdb/}.

Backbone ProToken embeddings for the PDB dataset is available on {http://ftp.cbi.pku.edu.cn/psp/true_structure_dataset/protoken/}.

A precompiled version of the ProToken model along with tutorials is available online via {https://colab.research.google.com/drive/15bBbfa7WigruoME089cSfE242K1MvRGz#scrollTo=vxQ7xhQQCBEc}, including the parameter checkpoints for the Encoder and Decoder as well as the Token vocabulary.

## Notes

### Competing Interest Statement

The authors have declared no competing interest.

### Summary of Updates

We added some new results and reorganized the sections in the manuscript.

## References

Abramson J, Adler J, Dunger J, Evans R, Green T, Pritzel A, Ronneberger O, Willmore L, Ballard AJ, Bambrick J, Bodenstein SW, Evans DA, Hung C-C, O’Neill M, Reiman D, Tunyasuvunakool K, Wu Z, Žemgulytė A, Arvaniti E, Beattie C, Bertolli O, Bridgland A, Cherepanov A, Congreve M, Cowen-Rivers AI, Cowie A, Figurnov M, Fuchs FB, Gladman H, Jain R, Khan YA, Low CMR, Perlin K, Potapenko A, Savy P, Singh S, Stecula A, Thillaisundaram A, Tong C, Yakneen S, Zhong ED, Zielinski M, Žídek A, Bapst V, Kohli P, Jaderberg M, Hassabis D, Jumper JM. 2024. Accurate structure prediction of biomolecular interactions with AlphaFold 3. Nature 630:493–500. doi:10.1038/s41586-024-07487-w

Alemi AA, Fischer I, Dillon JV, Murphy K. 2019. Deep Variational Information Bottleneck. doi:10.48550/arXiv.1612.00410

Banerjee A, Arora V. 2023. wav2tok: Deep Sequence Tokenizer for Audio RetrievalThe Eleventh International Conference on Learning Representations.

Basu S, Wallner B. 2016. DockQ: A Quality Measure for Protein-Protein Docking Models. PLOS ONE 11:e0161879. doi:10.1371/journal.pone.0161879

Bengio Y, Léonard N, Courville A. 2013. Estimating or Propagating Gradients Through Stochastic Neurons for Conditional Computation. doi:10.48550/arXiv.1308.3432

Berman HM, Westbrook J, Feng Z, Gilliland G, Bhat TN, Weissig H, Shindyalov IN, Bourne PE. 2000. The Protein Data Bank. Nucleic Acids Res 28:235–242. doi:10.1093/nar/28.1.235

Chang H, Zhang H, Jiang L, Liu C, Freeman WT. 2022. MaskGIT: Masked Generative Image Transformer. doi:10.48550/arXiv.2202.04200

Chen X, Duan Y, Houthooft R, Schulman J, Sutskever I, Abbeel P. 2016. InfoGAN: Interpretable Representation Learning by Information Maximizing Generative Adversarial Nets. doi:10.48550/arXiv.1606.03657

Chen X, Mishra N, Rohaninejad M, Abbeel P. 2017. PixelSNAIL: An Improved Autoregressive Generative Model. doi:10.48550/arXiv.1712.09763

del Alamo D, Sala D, Mchaourab HS, Meiler J. 2022. Sampling alternative conformational states of transporters and receptors with AlphaFold2. eLife 11:e75751. doi:10.7554/eLife.75751

Dosovitskiy A, Beyer L, Kolesnikov A, Weissenborn D, Zhai X, Unterthiner T, Dehghani M, Minderer M, Heigold G, Gelly S, Uszkoreit J, Houlsby N. 2021. An Image is Worth 16×16 Words: Transformers for Image Recognition at Scale. doi:10.48550/arXiv.2010.11929

Eaton WA, Muñoz V, Hagen SJ, Jas GS, Lapidus LJ, Henry ER, Hofrichter J. 2000. Fast Kinetics and Mechanisms in Protein Folding1. Annu Rev Biophys 29:327–359. doi:10.1146/annurev.biophys.29.1.327

Esser P, Rombach R, Ommer B. 2021. Taming Transformers for High-Resolution Image Synthesis. Presented at the Proceedings of the IEEE/CVF Conference on Computer Vision and Pattern Recognition. pp. 12873– 12883.

Gao M, Nakajima An D, Parks JM, Skolnick J. 2022. AF2Complex predicts direct physical interactions in multimeric proteins with deep learning. Nat Commun 13:1744. doi:10.1038/s41467-022-29394-2

Ghosh DK, Ranjan A. 2020. The metastable states of proteins. Protein Sci 29:1559–1568. doi:10.1002/pro.3859

Goodfellow I, Pouget-Abadie J, Mirza M, Xu B, Warde-Farley D, Ozair S, Courville A, Bengio Y. 2014. Generative Adversarial NetsAdvances in Neural Information Processing Systems. Curran Associates, Inc.

Goodfellow IJ, Shlens J, Szegedy C. 2015. Explaining and Harnessing Adversarial Examples. doi:10.48550/arXiv.1412.6572

Henikoff S, Henikoff JG. 1992. Amino acid substitution matrices from protein blocks. Proc Natl Acad Sci 89:10915–10919. doi:10.1073/pnas.89.22.10915

Ho J, Jain A, Abbeel P. 2020. Denoising Diffusion Probabilistic ModelsAdvances in Neural Information Processing Systems. Curran Associates, Inc. pp. 6840–6851.

Hrabe T, Li Z, Sedova M, Rotkiewicz P, Jaroszewski L, Godzik A. 2016. PDBFlex: exploring flexibility in protein structures. Nucleic Acids Res 44:D423–428. doi:10.1093/nar/gkv1316

Ingraham JB, Baranov M, Costello Z, Barber KW, Wang W, Ismail A, Frappier V, Lord DM, Ng-Thow-Hing C, Van Vlack ER, Tie S, Xue V, Cowles SC, Leung A, Rodrigues JV, Morales-Perez CL, Ayoub AM, Green R, Puentes K, Oplinger F, Panwar NV, Obermeyer F, Root AR, Beam AL, Poelwijk FJ, Grigoryan G. 2023. Illuminating protein space with a programmable generative model. Nature. doi:10.1038/s41586-023-06728-8

Jing B, Berger B, Jaakkola T. 2024. AlphaFold Meets Flow Matching for Generating Protein Ensembles. doi:10.48550/arXiv.2402.04845

Jing B, Erives E, Pao-Huang P, Corso G, Berger B, Jaakkola T. 2023. EigenFold: Generative Protein Structure Prediction with Diffusion Models. doi:10.48550/arXiv.2304.02198

Jumper J, Evans R, Pritzel A, Green T, Figurnov M, Ronneberger O, Tunyasuvunakool K, Bates R, Žídek A, Potapenko A, Bridgland A, Meyer C, Kohl SAA, Ballard AJ, Cowie A, Romera-Paredes B, Nikolov S, Jain R, Adler J, Back T, Petersen S, Reiman D, Clancy E, Zielinski M, Steinegger M, Pacholska M, Berghammer T, Bodenstein S, Silver D, Vinyals O, Senior AW, Kavukcuoglu K, Kohli P, Hassabis D. 2021. Highly accurate protein structure prediction with AlphaFold. Nature 596:583–589. doi:10.1038/s41586-021-03819-2

Kalakoti Y, Wallner B. 2024. AFsample2: Predicting multiple conformations and ensembles with AlphaFold2. doi:10.1101/2024.05.28.596195

Kingma DP, Welling M. 2022. Auto-Encoding Variational Bayes. doi:10.48550/arXiv.1312.6114

Konovalov KA, Unarta IC, Cao S, Goonetilleke EC, Huang X. 2021. Markov State Models to Study the Functional Dynamics of Proteins in the Wake of Machine Learning. JACS Au 1:1330–1341. doi:10.1021/jacsau.1c00254

Łańcucki A, Chorowski J, Sanchez G, Marxer R, Chen N, Dolfing Hjga, Khurana S, Alumäe T, Laurent A. 2020. Robust Training of Vector Quantized Bottleneck Models. doi:10.48550/arXiv.2005.08520

Li C-L, Chang W-C, Cheng Y, Yang Y, Póczos B. 2017. MMD GAN: Towards Deeper Understanding of Moment Matching Network. doi:10.48550/arXiv.1705.08584

Liu L, Dong C, Liu X, Yu B, Gao J. 2023. Bridging Discrete and Backpropagation: Straight-Through and Beyond. Adv Neural Inf Process Syst 36:12291–12311.

Liu S, Zhang J, Chu H, Wang M, Xue B, Ni N, Yu J, Xie Y, Chen Z, Chen M, Liu Y, Patra P, Xu F, Chen J, Wang Z, Yang L, Yu F, Chen L, Gao YQ. 2022. PSP: Million-level Protein Sequence Dataset for Protein Structure Prediction. doi:10.48550/arXiv.2206.12240

Liu Y, Zhang K, Li Y, Yan Z, Gao C, Chen R, Yuan Z, Huang Y, Sun H, Gao J, He L, Sun L. 2024. Sora: A Review on Background, Technology, Limitations, and Opportunities of Large Vision Models. doi:10.48550/arXiv.2402.17177

Ma P, Li D-W, Brüschweiler R. 2023. Predicting protein flexibility with AlphaFold. Proteins Struct Funct Bioinforma 91:847–855. doi:10.1002/prot.26471

Mariani V, Biasini M, Barbato A, Schwede T. 2013. lDDT: a local superposition-free score for comparing protein structures and models using distance difference tests. Bioinformatics 29:2722–2728. doi:10.1093/bioinformatics/btt473

Markov state models of biomolecular conformational dynamics. 2014. Curr Opin Struct Biol 25:135–144. doi:10.1016/j.sbi.2014.04.002

Mikolov T, Sutskever I, Chen K, Corrado G, Dean J. 2013. Distributed Representations of Words and Phrases and their Compositionality. doi:10.48550/arXiv.1310.4546

Mirza M, Osindero S. 2014. Conditional Generative Adversarial Nets. doi:10.48550/arXiv.1411.1784

Monticelli L, Kandasamy SK, Periole X, Larson RG, Tieleman DP, Marrink S-J. 2008. The MARTINI Coarse-Grained Force Field: Extension to Proteins. J Chem Theory Comput 4:819–834. doi:10.1021/ct700324x

Oord A van den, Vinyals O, Kavukcuoglu K. 2018. Neural Discrete Representation Learning. doi:10.48550/arXiv.1711.00937

OpenAI, Achiam J, Adler S, Agarwal S, Ahmad L, Akkaya I, Aleman FL, Almeida D, Altenschmidt J, Altman S, Anadkat S, Avila R, Babuschkin I, Balaji S, Balcom V, Baltescu P, Bao H, Bavarian M, Belgum J, Bello I, Berdine J, Bernadett-Shapiro G, Berner C, Bogdonoff L, Boiko O, Boyd M, Brakman A-L, Brockman G, Brooks T, Brundage M, Button K, Cai T, Campbell R, Cann A, Carey B, Carlson C, Carmichael R, Chan B, Chang C, Chantzis F, Chen D, Chen S, Chen R, Chen J, Chen M, Chess B, Cho C, Chu C, Chung HW, Cummings D, Currier J, Dai Y, Decareaux C, Degry T, Deutsch N, Deville D, Dhar A, Dohan D, Dowling S, Dunning S, Ecoffet A, Eleti A, Eloundou T, Farhi D, Fedus L, Felix N, Fishman SP, Forte J, Fulford I, Gao L, Georges E, Gibson C, Goel V, Gogineni T, Goh G, Gontijo-Lopes R, Gordon J, Grafstein M, Gray S, Greene R, Gross J, Gu SS, Guo Y, Hallacy C, Han J, Harris J, He Y, Heaton M, Heidecke J, Hesse C, Hickey A, Hickey W, Hoeschele P, Houghton B, Hsu K, Hu S, Hu X, Huizinga J, Jain Shantanu, Jain Shawn, Jang J, Jiang A, Jiang R, Jin H, Jin D, Jomoto S, Jonn B, Jun H, Kaftan T, Kaiser Ł, Kamali A, Kanitscheider I, Keskar NS, Khan T, Kilpatrick L, Kim JW, Kim C, Kim Y, Kirchner JH, Kiros J, Knight M, Kokotajlo D, Kondraciuk Ł, Kondrich A, Konstantinidis A, Kosic K, Krueger G, Kuo V, Lampe M, Lan I, Lee T, Leike J, Leung J, Levy D, Li CM, Lim R, Lin M, Lin S, Litwin M, Lopez T, Lowe R, Lue P, Makanju A, Malfacini K, Manning S, Markov T, Markovski Y, Martin B, Mayer K, Mayne A, McGrew B, McKinney SM, McLeavey C, McMillan P, McNeil J, Medina D, Mehta A, Menick J, Metz L, Mishchenko A, Mishkin P, Monaco V, Morikawa E, Mossing D, Mu T, Murati M, Murk O, Mély D, Nair A, Nakano R, Nayak R, Neelakantan A, Ngo R, Noh H, Ouyang L, O’Keefe C, Pachocki J, Paino A, Palermo J, Pantuliano A, Parascandolo G, Parish J, Parparita E, Passos A, Pavlov M, Peng A, Perelman A, Peres F de AB, Petrov M, Pinto HP de O, Michael, Pokorny, Pokrass M, Pong VH, Powell T, Power A, Power B, Proehl E, Puri R, Radford A, Rae J, Ramesh A, Raymond C, Real F, Rimbach K, Ross C, Rotsted B, Roussez H, Ryder N, Saltarelli M, Sanders T, Santurkar S, Sastry G, Schmidt H, Schnurr D, Schulman J, Selsam D, Sheppard K, Sherbakov T, Shieh J, Shoker S, Shyam P, Sidor S, Sigler E, Simens M, Sitkin J, Slama K, Sohl I, Sokolowsky B, Song Y, Staudacher N, Such FP, Summers N, Sutskever I, Tang J, Tezak N, Thompson MB, Tillet P, Tootoonchian A, Tseng E, Tuggle P, Turley N, Tworek J, Uribe JFC, Vallone A, Vijayvergiya A, Voss C, Wainwright C, Wang JJ, Wang A, Wang B, Ward J, Wei J, Weinmann CJ, Welihinda A, Welinder P, Weng J, Weng L, Wiethoff M, Willner D, Winter C, Wolrich S, Wong H, Workman L, Wu S, Wu J, Wu M, Xiao K, Xu T, Yoo S, Yu K, Yuan Q, Zaremba W, Zellers R, Zhang C, Zhang M, Zhao S, Zheng T, Zhuang J, Zhuk W, Zoph B. 2024. GPT-4 Technical Report. doi:10.48550/arXiv.2303.08774openmm/pdbfixer. 2024.

Papamakarios G, Pavlakou T, Murray I. 2017. Masked Autoregressive Flow for Density EstimationAdvances in Neural Information Processing Systems. Curran Associates, Inc.

Peebles W, Xie S. 2023. Scalable Diffusion Models with Transformers. doi:10.48550/arXiv.2212.09748

Rombach R, Blattmann A, Lorenz D, Esser P, Ommer B. 2022. High-Resolution Image Synthesis With Latent Diffusion Models. Presented at the Proceedings of the IEEE/CVF Conference on Computer Vision and Pattern Recognition. pp. 10684–10695.

Ryoo M, Piergiovanni A, Arnab A, Dehghani M, Angelova A. 2021. TokenLearner: Adaptive Space-Time Tokenization for Videos In: Ranzato M, Beygelzimer A, Dauphin Y, Liang PS, Vaughan JW, editors. Advances in Neural Information Processing Systems. Curran Associates, Inc. pp. 12786–12797.

Salimans T, Karpathy A, Chen X, Kingma DP. 2017. PixelCNN++: Improving the PixelCNN with Discretized Logistic Mixture Likelihood and Other Modifications. doi:10.48550/arXiv.1701.05517

Senior AW, Evans R, Jumper J, Kirkpatrick J, Sifre L, Green T, Qin C, Žídek A, Nelson AWR, Bridgland A, Penedones H, Petersen S, Simonyan K, Crossan S, Kohli P, Jones DT, Silver D, Kavukcuoglu K, Hassabis 2020. Improved protein structure prediction using potentials from deep learning. Nature 577:706–710. doi:10.1038/s41586-019-1923-7

Song Y, Ermon S. 2019. Generative Modeling by Estimating Gradients of the Data Distribution. Adv Neural Inf Process Syst 32.

Song Y, Sohl-Dickstein J, Kingma DP, Kumar A, Ermon S, Poole B. 2021. Score-Based Generative Modeling through Stochastic Differential Equations. doi:10.48550/arXiv.2011.13456

Sun P, Jiang Y, Chen S, Zhang S, Peng B, Luo P, Yuan Z. 2024. Autoregressive Model Beats Diffusion: Llama for Scalable Image Generation. doi:10.48550/arXiv.2406.06525

Sun Z, Liu Q, Qu G, Feng Y, Reetz MT. 2019. Utility of B-Factors in Protein Science: Interpreting Rigidity, Flexibility, and Internal Motion and Engineering Thermostability. Chem Rev 119:1626–1665. doi:10.1021/acs.chemrev.8b00290

Taketomi H, Ueda Y, Gō N. 1975. Studies on protein folding, unfolding and fluctuations by computer simulation. The effect of specific amino acid sequence represented by specific inter-unit interactions. Int J Pept Protein Res 7:445–459.

Tian K, Jiang Y, Yuan Z, Peng B, Wang L. 2024. Visual Autoregressive Modeling: Scalable Image Generation via Next-Scale Prediction. doi:10.48550/arXiv.2404.02905

van Kempen M, Kim SS, Tumescheit C, Mirdita M, Lee J, Gilchrist CLM, Söding J, Steinegger M. 2024. Fast and accurate protein structure search with Foldseek. Nat Biotechnol 42:243–246. doi:10.1038/s41587-023-01773-0

Varadi M, Bertoni D, Magana P, Paramval U, Pidruchna I, Radhakrishnan M, Tsenkov M, Nair S, Mirdita M, Yeo J, Kovalevskiy O, Tunyasuvunakool K, Laydon A, Žídek A, Tomlinson H, Hariharan D, Abrahamson J, Green T, Jumper J, Birney E, Steinegger M, Hassabis D, Velankar S. 2024. AlphaFold Protein Structure Database in 2024: providing structure coverage for over 214 million protein sequences. Nucleic Acids Res 52:D368–D375. doi:10.1093/nar/gkad1011

Wales D. 2004. Energy Landscapes: Applications to Clusters, Biomolecules and Glasses, Cambridge Molecular Science. Cambridge: Cambridge University Press. doi:10.1017/CBO9780511721724

Watson JL, Juergens D, Bennett NR, Trippe BL, Yim J, Eisenach HE, Ahern W, Borst AJ, Ragotte RJ, Milles LF, Wicky BIM, Hanikel N, Pellock SJ, Courbet A, Sheffler W, Wang J, Venkatesh P, Sappington I, Torres SV, Lauko A, De Bortoli V, Mathieu E, Ovchinnikov S, Barzilay R, Jaakkola TS, DiMaio F, Baek M, Baker D. 2023. De novo design of protein structure and function with RFdiffusion. Nature 620:1089–1100. doi:10.1038/s41586-023-06415-8

Wayment-Steele HK, Ojoawo A, Otten R, Apitz JM, Pitsawong W, Hömberger M, Ovchinnikov S, Colwell L, Kern D. 2024. Predicting multiple conformations via sequence clustering and AlphaFold2. Nature 625:832–839. doi:10.1038/s41586-023-06832-9

Yu J, Li X, Koh JY, Zhang H, Pang R, Qin J, Ku A, Xu Y, Baldridge J, Wu Y. 2022. Vector-quantized Image Modeling with Improved VQGAN. doi:10.48550/arXiv.2110.04627

Zhang J, Lei Y-K, Che X, Zhang Z, Yang YI, Gao YQ. 2019. Deep Representation Learning for Complex Free-Energy Landscapes. J Phys Chem Lett. doi:10.1021/acs.jpclett.9b02012

Zhang J, Lin X E W, Gao YQ. 2023a. Machine-Learned Invertible Coarse Graining for Multiscale Molecular Modeling. doi:10.48550/arXiv.2305.01243

Zhang J, Liu S, Chen M, Chu H, Wang M, Wang Z, Yu J, Ni N, Yu F, Chen D, Yang YI, Xue B, Yang L, Liu Y, Gao YQ. 2023b. Unsupervisedly Prompting AlphaFold2 for Accurate Few-Shot Protein Structure Prediction. J Chem Theory Comput. doi:10.1021/acs.jctc.3c00528

