## Supplementary Information for "Tokenizing Foldable Protein Structures with Machine-Learned Artificial Amino-Acid Vocabulary"

---

#### Table of Contents

Supplementary Figures
Figure S1
Figure S2
Figure S3

Supplementary Notes
Note S1. Model Details
Note S2. Optimization Details
Note S3. Data Details

Reference

### Supplementary Figures

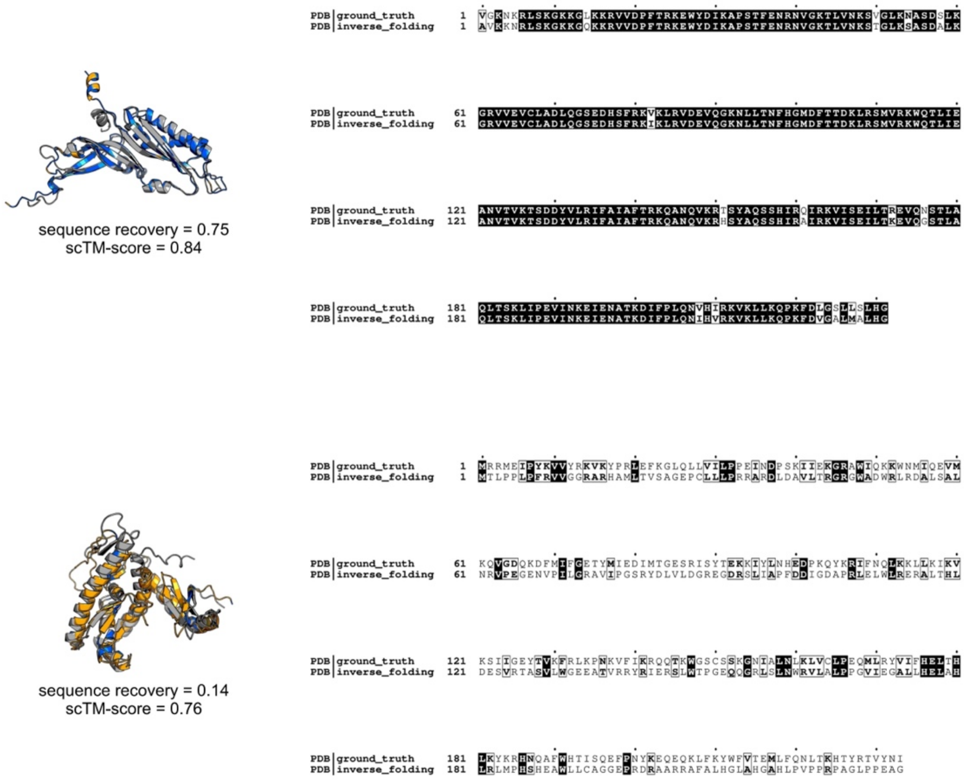


**Figure S1. Case Study of Inverse Folding Tasks empowered with PT-DiT.** The ground truth structure (gray) and structure predicted by ESMFold based on the generated sequence of PT-DiT are aligned and shown on the left. The common residues are colored in marine and uncommon residues in orange. The ground truth sequence and sequence generated by PT-DiT are aligned on the right. Tshe common residues are colored in black background. The upper case has a high sequence recovery and high self-consistency TM-score of 0.84, while the lower case has a low sequence recovery, while maintain relatively high self-consistency TM-score of 0.76.

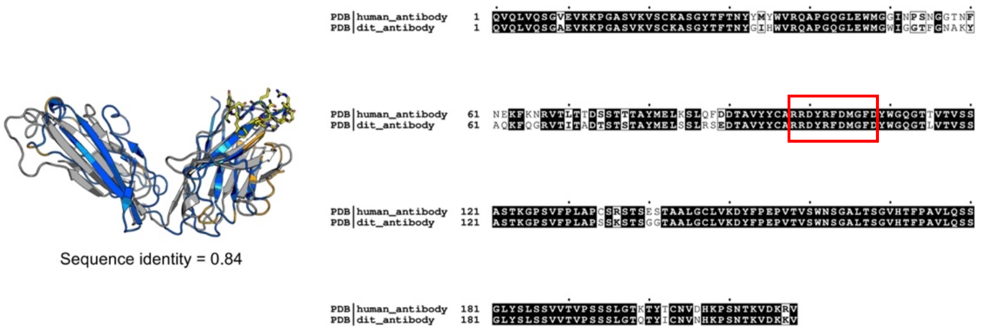


**Figure S2. Comparison of sequence and structure between PT-DiT generated antibody candidates and a human antibody.** The sequence generated by PT-DiT compared to the human antibody sequence. The CDR3 residues are marked in red square and the common residues in black background. Sequence identity of two structures is 0.84.

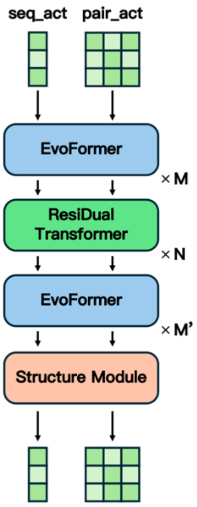


**Figure S3. Model Structure of Structure Encoder.** Single activation and pair activation are sent to the 'sandwich' hyperformer, which consists of M layer of EvoFormer module and N layer of ResiDual Transformer module. Then the outputs go through the M' layer of Structure Module and get the updated single activation and pair activation.

### Supplementary Notes

#### Note S1. Model Details

##### A. Decoder

The Decoder is a composite function consisting of a *Token Duplicator* module and a *Detokenizer* module.

###### 1. Detokenizer

The Detokenizer module is a SE(3)-equivariant generative model which samples protein structures from a metastable ensemble corresponding to a given ProToken string. More specifically, the all-atom structures are generated in a factorized way by a backbone detokenizer $g_{\phi_1}$ and a sidechain detokenizer $g_{\phi_2}$, respectively,

$$\mathbf{x} \sim p_{\phi}(\mathbf{x}_{\mathrm{BB}}, \mathbf{x}_{\mathrm{SC}}) = p_{\phi_1}(\mathbf{x}_{\mathrm{BB}}) p_{\phi_2}(\mathbf{x}_{\mathrm{SC}} | \mathbf{x}_{\mathrm{BB}}) \tag{S1}$$

$$p_{\phi_1}(\mathbf{x}_{\mathrm{BB}}) \mathrm{d}\mathbf{x}_{\mathrm{BB}} = g_{\phi_1}(\boldsymbol{\epsilon}; \mathbf{z}_{\mathrm{BB}})) \mathrm{d}\boldsymbol{\epsilon} \tag{S2}$$

$$p_{\phi_2}(\mathbf{x}_{\mathrm{SC}} | \mathbf{x}_{\mathrm{BB}}) \mathrm{d}\mathbf{x}_{\mathrm{SC}} = g_{\phi_2}(\boldsymbol{\epsilon}; \mathbf{z}_{\mathrm{SC}}, \mathbf{x}_{\mathrm{BB}}) \mathrm{d}\boldsymbol{\epsilon} \tag{S3}$$

where the backbone structure is first generated according to the $p_{\phi}(\mathbf{x}_{\mathrm{BB}})$ conditioned on the backbone tokens $\mathbf{z}_{\mathrm{BB}}$, followed by the generation of sidechains from $p(\mathbf{x}_{\mathrm{SC}} | \mathbf{x}_{\mathrm{BB}})$ conditioned on the backbone structure $\mathbf{x}_{\mathrm{BB}}$ as well as the sidechain tokens $\mathbf{z}_{\mathrm{SC}}$.

Noteworthy, any unconditional backbone generative model (such as RFDiffusion(Watson et al., 2023)) can be adopted as an initializer for the backbone detokenizer further optimized through conditional fine-tuning. Similarly, any generative sidechain packer can be directly plugged-in (or fine-tuned) as the sidechain detokenizer. We implemented DLPacker(Misiura et al., 2022) in this work and did not fine-tune it.

###### 2. Token Duplicator

The duplicate set defined in Eq. 2 depends both on the volume of the ProToken space and the over-capacity of Decoder. The duplicate set associated with a less compact ProToken space, or a less invertible Decoder, intuitively tends to be larger, which is harmful to protein structure generation tasks as will be shown later.

Although the exact likelihood (Eq. 2) is hard to compute in practice, fortunately, we can develop proper lower bounds that can be conveniently used for maximum likelihood estimation of $\mathbf{x}$ through the transformed representation $\mathbf{z}$.

The first lower bound ($\mathcal{L}_1$) can be derived via the *truncation trick*. Given a (truncated) subset $\widetilde{\mathbb{Z}}(\mathbf{x}) \subseteq \mathbb{Z}(\mathbf{x}; \phi)$,

$$\log p(\mathbf{x}) = \log \sum_{\mathbf{z}\in\mathbb{Z}(\mathbf{x};\phi)} p(\mathbf{z}_i) \geq \log \sum_{\mathbf{z}\in\tilde{\mathbb{Z}}(\mathbf{x})} p(\mathbf{z}_i) := \mathcal{L}_1\left(\tilde{\mathbb{Z}}(\mathbf{x})\right) \tag{S4}$$

$\mathcal{L}_1$ exhibits an important property that,

$$\text{If } \tilde{\mathbb{Z}}_2(\mathbf{x}) \supset \tilde{\mathbb{Z}}_1(\mathbf{x}), \text{then } \mathcal{L}_1\left(\tilde{\mathbb{Z}}_2(\mathbf{x})\right) \geq \mathcal{L}_1\left(\tilde{\mathbb{Z}}_1(\mathbf{x})\right) \tag{S5}$$

The equality holds if elements in the difference set, $\tilde{\mathbb{Z}}_2(\mathbf{x}) - \tilde{\mathbb{Z}}_1(\mathbf{x})$, have zero probability density in total. $\mathcal{L}_1$ is still inconvenient to compute in practice because the summation of probability is prior to logrithm. Thus, we further develop another lower bound ($\mathcal{L}_2$) based on $\mathcal{L}_1$ according to Jensen's inequality,

$$\mathcal{L}_1\left(\tilde{\mathbb{Z}}(\mathbf{x})\right) = \log \frac{\sum_{\mathbf{z}\in\tilde{\mathbb{Z}}(\mathbf{x})} p(\mathbf{z}_i)}{\left|\tilde{\mathbb{Z}}(\mathbf{x})\right|} + \log\left|\tilde{\mathbb{Z}}(\mathbf{x})\right| \geq \frac{\sum_{\mathbf{z}\in\tilde{\mathbb{Z}}(\mathbf{x})} \log p(\mathbf{z}_i)}{\left|\tilde{\mathbb{Z}}(\mathbf{x})\right|} + \log\left|\tilde{\mathbb{Z}}(\mathbf{x})\right| \tag{S6}$$

$$\log p(\mathbf{x}) \geq \mathcal{L}_1\left(\tilde{\mathbb{Z}}(\mathbf{x})\right) \geq \mathbb{E}_{z\sim\tilde{p}(\mathbf{z}|\mathbf{x},\phi)} \log p(\mathbf{z}) + \log\left|\tilde{\mathbb{Z}}(\mathbf{x})\right| := \mathcal{L}_2\left(\tilde{\mathbb{Z}}(\mathbf{x})\right) \tag{S7}$$

Different from $\mathcal{L}_1$, Eq. S7 allows us to perform minibatch optimization by unbiasedly sampling from a $\tilde{\mathbb{Z}}(\mathbf{x})$ during training. Similarly, it also follows straightforwardly from Eq. S7 that the larger $\left|\tilde{\mathbb{Z}}(\mathbf{x})\right|$ is, the tighter the lower bound $\mathcal{L}_2$ is, indicating the importance of expanding the subset $\tilde{\mathbb{Z}}(\mathbf{x})$.

The remaining issue is how to construct and sample from the duplicate (sub-)set. A naïve approach is to approximate the duplicate set by a singleton set, i.e., $\tilde{\mathbb{Z}}(\mathbf{x}) = \{f_\theta(\mathbf{x})\}$, which leads to a suboptimal truncation for $\mathcal{L}_1$. Or one can determistically set $\mathbf{z} = f_\theta(\mathbf{x})$ and estimate the expectation term in $\mathcal{L}_2$, which leads to a highly biased and high-variance one-sample Monte Carlo estimator.

To circumvent these drawbacks, a *Token Duplicator* module, $q_\phi(\mathbf{z}|\mathbf{x}, g_\phi)$, is prepended to the Detokenizer, which is responsible to expand and sample from the duplicate subset $\tilde{\mathbb{Z}}(\mathbf{x})$. In general, any conditional generative model which yields diverse ProTokens which can be decoded back to $\mathbf{x}$ via the Decoder can be used as $q_\phi(\mathbf{z}|\mathbf{x}, g_\phi)$.

In this work, given a structure $\mathbf{x}$, we initialize the duplicate set with $\{\mathbf{z} = f_\theta(\mathbf{x})\}$, and implement a sampling-based, Monte-Carlo-style *Token Duplicator* to expanding the duplicate set. We randomly mutate the residue-wise backbone tokens in $\mathbf{z}$ according to the similarity matrix of token embeddings, yielding a perturbed $\mathbf{z}'$. We accept the proposed mutation $\mathbf{z}'$ with a probability proportional to the reconstruction quality of $g_\phi(\mathbf{z}')$, based on which the next residue-wise mutation is performed. If $g_\phi(\mathbf{z}')$ is sufficiently close to $\mathbf{x}$ (defined as TM-score($\mathbf{x}$, $g_\phi(\mathbf{z}')$) > 0.9), we add it to the duplicate set.

During generative training of protein structures $\mathbf{x}$ via ProTokens $\mathbf{z}$, we first resort to the *Token Duplicator* and construct a sufficiently large duplicate subset $\widetilde{\mathbb{Z}}(\mathbf{x})$. The $\mathcal{L}_1$ or $\mathcal{L}_2$ is then optimized based on $\widetilde{\mathbb{Z}}(\mathbf{x})$ in order to perform maximum likelihood estimation of $\mathbf{x}$. This approach is similar to data (or label) augmentation which plays key role in many state-of-the-art generative AI models.

#### B. Encoder

##### 1. Structure Encoder

Considering the separation of timescales, we encode the sidechain and backbone structures separately as in Eq. 4. As for the *backbone structure encoder* $f_{\theta_1}(\mathbf{x}_{\mathrm{BB}})$, an SE(3)-invariant Transformer based on invariant point attention(Jumper et al., 2021), is designed to transform the Cartesian coordinates of the N, CA, C, O atoms of each residue to a SE(3)-invariant vector. The model is composed of an SE(3)-invariant HyperFormer(Zhang et al., 2023) and a structure-aware Transformer (Figure S3). Specifically, the HyperFormer treats residues and inter-residue geometries as vertices and edges of a graph, respectively. Like EvoFormer in AlphaFold2, HyperFormer updates both the vertex and edge representations interdependently via hyper-attention mechanism. Based on the refined vertex (or single) and edge (or pair) representations output by HyperFormer, invariant point attention introduced in AlphaFold2 is adopted in the structure-aware Transformer, in order to efficiently learn the global geometric features (especially, long-range interactions) of a protein backbone.

On the other hand, since the equilibration of sidechain conformations is much faster than the backbone, all relaxed conformations of a sidechain (e.g., according to Boltzmann distribution) in the context of a given backbone can be considered as a single metastable state and reasonably embedded by a single vector which can distinguish the chemical identity of the sidechain fragment. Therefore, we transact the amino acid embedding learned by AF2 as the s*idechain structure encoder* $f_{\theta_2}(\mathbf{x}_{\mathrm{SC}})$ without further optimization.

The embedding of an all-atom structure is obtained via the Cartesian product operation, that is, by concatenating the backbone embedding and sidechain embedding.

##### 2. Deduplicator

Note that our choice of sidechain encoding is naturally invariant to perturbations of sidechain conformations. However, unlike the *sidechain structure encoder*, one particular concern of a learned *backbone structure encoder* is that the *robustness* of the yielded codes against subtle structure perturbation is not guaranteed. To alleviate this issue, we append a *Deduplicator* $u_\theta$ to the *backbone structure encoder* (Fig. 2c), which is trained to surjectively map structures which belong to the same metastable states (or merely differ mutually up to negligible perturbations) to almost the same embedding,

$$u_{\theta_1} \circ f_{\theta_1}(\mathbf{x}_{\mathrm{BB}}) \approx u_{\theta_1} \circ f_{\theta_1}(\mathbf{x}_{\mathrm{BB}} + \Delta\mathbf{x}_{\mathrm{BB}}) \tag{S8}$$

where $\Delta\mathbf{x}_{\mathrm{BB}}$ denotes the structural fluctuation within a metastable state. In terms of physics, the *Deduplicator* behaves like a "relaxation simulator" which relaxes fluctuated structures within a metastable ensemble towards a single stable representative structure, and consequently, degenerates the embeddings for fluctuated structures.

### C. Tokenizer

The Tokenizer is a composition of a *Clustering* module $s_\theta$ (which has been elaborated at length in the main text), and a *Compressing* module $r_\theta$.

#### 1. Compressing Module

The ProToken of all-atom structure is obtained via $\mathbf{z} = \mathbf{z}_{\mathrm{BB}} \otimes \mathbf{z}_{\mathrm{SC}}$, that is, the Cartesian product of the backbone token set and the sidechain token set. The continous ProToken embedding is equivalent to concatenating the the backbone embedding and sidechain embedding together, which has the shape of $(N_{\mathrm{res}}, d = d_{\mathrm{BB}} + d_{\mathrm{SC}})$ for a $N_{\mathrm{res}}$-long protein.

Note that without processing, $d_{\mathrm{BB}}$ and $d_{\mathrm{SC}}$ are usually large such that $d \gg 3\bar{N}_{\mathrm{atom}}$ where $\bar{N}_{\mathrm{atom}}$ stands for the average number of atoms per residue. For instance, in our model, $d_{\mathrm{BB}} = 32$ during training and $d_{\mathrm{SC}} = 256$ in consistency with AF2. Despite of being SE(3)-invariant, the dimensionality of such a representation is still much too higher than the intrinsic degrees of freedoms of a protein (which is upper bounded by the number of Cartesian coordinates of its structure).

Therefore, we include a *Compressing* module $r_\theta(\mathbf{v})\colon \mathbb{R}^{N_{\mathrm{res}}*d} \to \mathbb{R}^{Q*c}$ to concentrating the information of ProToken by lowering its dimensionality. The compression can be performed lengthwise and (or) depth-wise. As shown in Fig. 2d, for the lengthwise compression, we transformed a ProToken string of shape $(N_{\mathrm{res}}, d)$ into $(Q, d)$, with $Q$ being a predefined number independent of and usaually smaller than the average $N_{\mathrm{res}}$. In this research, we performed the depth-wise compression (Fig. 2d), which reduces the shape of $(N_{\mathrm{res}}, d)$ into $(N_{\mathrm{res}}, c \ll d)$ , and we achieved this goal by means of dimensionality reduction methods(McInnes et al., 2020).

### Note S2. Optimization Details

#### A. ProToken Distiller

##### 1. Training settings

ProToken Distiller is trained with a batch size of 288. The learning rate is set to 5e-4 with a cosine decay down to 2e-5 after 80,000 steps. The training is executed on 48 NVIDIA A100 GPUs.

The training is split into two stages. For the first 100,000 steps, we implemented the robustness loss $L_{\mathrm{ROB}}$ in Eq. 14, with a prepared set of adversarial examples but turned off the mutual information loss $L_{\mathrm{MI}}$ in Eq. 12. For the remaining 100,000 steps, we switched off the robustness loss, instead, MI loss was turned on.

##### 2. Metastable Perturbation Sampling (MPS)

To sample more metastable conformations of a 3D structure $\mathbf{x}_0$ associated with the same function, we performed metastable perturbation sampling according to the given 3D structure and yield $\{\mathbf{x}\}$. Specifically, we recommend several options that can serve for MPS: 1) resorting to a MD simulation engine and running temporal proximal sampling(Zhang et al., 2019); 2) resorting to AI-based sampler which can yield perturbed conformations like AF-Cluster(Wayment-Steele et al., 2024); 3) self-distillation of a pre-trained probabilistic ProToken distiller. In this research, we implemented the first two strategies to obtain perturbed metastable conformations corresponding to a given reference 3D structure.

#### B. ProToken Diffusion Transformer (PT-DiT)

To prepare the training data of PT-DiT, we first performed Encoder inference for the training set of protein structures, yielding a basic set of ProTokens containing 551957 ProToken samples.

In order to counteract the biased estimation of likelihood of the latent diffusion model, we augmented the basic ProToken set with the duplicate set obtained by the *Token Duplicator*. Through experiments, we found empirically that augmentation of latent duplicates is vital for the success of training PT-DiT.

The generation quality of PT-DiT can be susceptible to errors that cause “token switch”, so we introduced anisotropic diffusion kernel(Lin et al., 2024) for the variance-preserving diffusion process, in order to better align with the objective of the latent diffusion model

We trained PT-DiT with a batch size of 256, learning rate of 2e-4 for 1,000,000 steps, on 8 NVIDIA A100 GPUs.

### Note S3. Data Details

#### A. ProToken Distiller Training Dataset

Based on single-chain protein structures obtained from the RCSB Protein Data Bank (PDB) released before October 13, 2021, we performed data cleaning and filtering. We retained chains without structural gaps, excluded structures shorter than 30 residues, and omitted those derived from NMR experiments due to potential metastability issues. Ultimately, 551957 single-chain protein structures were used as training data. The list of PDB IDs and chain IDs is publicly accessible in the Open Science Framework (OSF) repository(Lin et al., 2023). These PDB structures are accessible from the RCSB website (www.rcsb.org) using the corresponding PDB and chain IDs.

#### B. ProToken Distiller Test Dataset

##### 1. Validation dataset

We used the validation dataset from the PSP dataset(Liu et al., 2022) for validation of single chain reconstruction task. It includes two main parts: the CASP14 dataset and a new validation dataset. The new dataset was curated from two source of public data. One is CAMEO targets from October 16, 2021, to February 12, 2022, and another is new single clusters from the PDB with 40% identity between October 13, 2021, and March 15, 2022.

Thus, the validation dataset is unique and diverse compared to the training set. We filtered out samples shorter than 1536 residues for easier validation. In total, we have 513 non-overlapping samples. All ground truth structures were released after October 13, 2021, which is after all the training samples. This prevents any data leak during validation and testing. The validation dataset is provided in the supplementary data.

##### 2. CASP14 single domain dataset

To clearly measure the performance of ProToken Distiller on the widely recognized CASP14 dataset, we included all 87 single-domain regular targets with ground truth structures in CASP14 to form this dataset, even though this dataset is part of the earlier validation set. The CASP14 single-domain structures are available in the OSF repository(Lin et al., 2023).

##### 3. CASP15 single domain dataset

All regular targets in CASP15 with ground truth structures are used to form this dataset, which results in 45 single domain protein structures. the CASP15 single domain dataset is available in the OSF repository(Lin et al., 2023).

##### 4. AFDB dark cluster dataset

Foldseek has identified 711,705 dark clusters, which are likely enriched with novel structures(“Clustering predicted structures at the scale of the known protein universe | Nature,” n.d.). To ensure structural quality, we followed Foldseek data processing flow and selected 33,842 clusters with the highest average AlphaFold2 prediction confidence (average plDDT >90). From each cluster, we chose the member with the highest confidence for further investigation, similar to Foldseek. The structures in this dark cluster dataset can be downloaded from the AlphaFoldDB website (https://alphafold.ebi.ac.uk/), and the name list is available at (https://afdb-cluster.steineggerlab.workers.dev/).

#### 5. CASP14 and CASP15 multi-domain dataset

CASP14 and CASP15 have provided "Domain Definitions" on their official website, which are used for multi-domain structure prediction tasks. Following the curation approach of DeepAssembly(“Multi-domain and complex protein structure prediction using inter-domain interactions from deep learning | Communications Biology,” n.d.), we formed a total of 30 multi-domain targets: 17 from CASP14 and 13 from CASP15. These targets were used to evaluate the reconstruction ability for multi-domain protein structures. The 30 multi-domain structures are available at the OSF repository(Lin et al., 2023).

#### 6. Multimer dataset

Datasets for the test of multi-chain protein reconstruction task were curated from AF2Complex benchmark sets(Gao et al., 2022), specifically Dimer1193 and Oligomer562. Since no multimer structures were used during the training of ProToken Distiller, all multimer cases in these two sets were reserved for benchmarking the multimer reconstruction task.

#### 7. PDBFlex conformation dataset

The PDBFlex database has organized structure clusters with similar sequences but significant structural differences(Hrabe et al., 2016). To assess ProToken's ability to distinguish and reconstruct various metastable states, we followed the database's definitions of Local RMSD and categorized the clusters into bins based on Local RMSD ranges of 2-4Å, 4-8Å, 8-16Å, 16-32Å, and 32-64Å. From each bin, we selected 10 clusters, totaling 50 structure clusters. From each cluster, we chose the pair of structures with the highest backbone RMSD, creating a final dataset of 100 structures. This dataset serves as the test set for the multi-conformation reconstruction task and all structures are available at the OSF repository(Lin et al., 2023).

#### 8. APObind Pocket-Ligand Dataset

Based on the APObind ligand unbound protein conformations(Aggarwal et al., 2021), we curated 229 *apo* structures aligned with their corresponding *holo* protein-ligand structures. In total, 458 proteins were used for the pocket reconstruction test. The pocket is defined according to the AF3 methodology, encompassing all heavy atoms within 10 Å of any heavy atom of the ligand. The backbone of a residue includes the ‘N’, ‘C’, ‘CA’, and ‘O’ atoms. The pocket backbone RMSD was calculated after aligning all backbone atoms of the *apo-holo* conformer pairs using Biopython and Numpy in Python scripts. The pocket residue indexes, *apo-holo* protein structure pairs together with the ligand structures are available at the OSF repository(Lin et al., 2023).

##### 9. Antibody CDR dataset

Following the DeepAb test set("Antibody structure prediction using interpretable deep learning," 2022), we used 92 antibody cases from the RosettaAntibody benchmark set (47 targets) and a set of clinical-stage therapeutic antibodies (45 targets) to form the antibody CDR dataset. We cleaned and annotated these protein complexes, resulting in a total of 238 single-chain structures. The Chothia CDR loop definitions were used to measure RMSD throughout this work. All the single-chain structures, FASTAs and annotations are available at the OSF repository(Lin et al., 2023).

#### C. PT-DiT Training Dataset

We randomly utilized 550957 ProToken sequences of the structures and corresponding amino acid sequences from the ProToken Distiller training set for PT-DiT training and the rest 1,000 data points serve as the validation set. The name lists for both the training and validation sets are available at the OSF repository(Lin et al., 2023).

### Reference


Aggarwal R, Gupta A, Priyakumar UD. 2021. APObind: A Dataset of Ligand Unbound Protein Conformations for Machine Learning Applications in De Novo Drug Design. doi:10.48550/arXiv.2108.09926

Antibody structure prediction using interpretable deep learning. 2022. . *Patterns* **3**:100406. doi:10.1016/j.patter.2021.100406

Clustering predicted structures at the scale of the known protein universe | Nature. n.d. https://www.nature.com/articles/s41586-023-06510-w

Gao M, Nakajima An D, Parks JM, Skolnick J. 2022. AF2Complex predicts direct physical interactions in multimeric proteins with deep learning. *Nat Commun* **13**:1744. doi:10.1038/s41467-022-29394-2

Hrabe T, Li Z, Sedova M, Rotkiewicz P, Jaroszewski L, Godzik A. 2016. PDBFlex: exploring flexibility in protein structures. *Nucleic Acids Res* **44**:D423-428. doi:10.1093/nar/gkv1316

Jumper J, Evans R, Pritzel A, Green T, Figurnov M, Ronneberger O, Tunyasuvunakool K, Bates R, Žídek A, Potapenko A, Bridgland A, Meyer C, Kohl SAA, Ballard AJ, Cowie A, Romera-Paredes B, Nikolov S, Jain R, Adler J, Back T, Petersen S, Reiman D, Clancy E, Zielinski M, Steinegger M, Pacholska M, Berghammer T, Bodenstein S, Silver D, Vinyals O, Senior AW, Kavukcuoglu K, Kohli P, Hassabis D. 2021. Highly accurate protein structure prediction with AlphaFold. *Nature* **596**:583–589.

doi:10.1038/s41586-021-03819-2

Lin X, Chen Z, Li Y. 2023. ProToken [Data & Source Code].

Lin X, Xia Y, Huang Y, Liu S, Zhang J, Gao YQ. 2024. Versatile Molecular Editing via Multimodal and Group-optimized Generative Learning. doi:10.26434/chemrxiv-2023-j2n6l-v2

Liu S, Zhang J, Chu H, Wang M, Xue B, Ni N, Yu J, Xie Y, Chen Z, Chen M, Liu Y, Patra P, Xu F, Chen J, Wang Z, Yang L, Yu F, Chen L, Gao YQ. 2022. PSP: Million-level Protein Sequence Dataset for Protein Structure Prediction. doi:10.48550/arXiv.2206.12240

McInnes L, Healy J, Melville J. 2020. UMAP: Uniform Manifold Approximation and Projection for Dimension Reduction. doi:10.48550/arXiv.1802.03426

Misiura M, Shroff R, Thyer R, Kolomeisky AB. 2022. DLPacker: Deep learning for prediction of amino acid side chain conformations in proteins. *Proteins Struct Funct Bioinforma* **90**:1278–1290. doi:10.1002/prot.26311

Multi-domain and complex protein structure prediction using inter-domain interactions from deep learning | Communications Biology. n.d. https://www.nature.com/articles/s42003-023-05610-7

Watson JL, Juergens D, Bennett NR, Trippe BL, Yim J, Eisenach HE, Ahern W, Borst AJ, Ragotte RJ, Milles LF, Wicky BIM, Hanikel N, Pellock SJ, Courbet A, Sheffler W, Wang J, Venkatesh P, Sappington I, Torres SV, Lauko A, De Bortoli V, Mathieu E, Ovchinnikov S, Barzilay R, Jaakkola TS, DiMaio F, Baek M, Baker D. 2023. De novo design of protein structure and function with RFdiffusion. *Nature* **620**:1089–1100. doi:10.1038/s41586-023-06415-8

Wayment-Steele HK, Ojoawo A, Otten R, Apitz JM, Pitsawong W, Hömberger M, Ovchinnikov S, Colwell L, Kern D. 2024. Predicting multiple conformations via sequence clustering and AlphaFold2. *Nature* **625**:832–839. doi:10.1038/s41586-023-06832-9

Zhang J, Lei Y-K, Che X, Zhang Z, Yang YI, Gao YQ. 2019. Deep Representation Learning for Complex Free-Energy Landscapes. *J Phys Chem Lett*. doi:10.1021/acs.jpclett.9b02012

Zhang J, Lei Y-K, Zhou Y, Yang YI, Gao YQ. 2023. Molecular CT: Unifying Geometry and Representation Learning for Molecules at Different Scales. doi:10.48550/arXiv.2012.11816
